# Usherin in the pineal gland: altered sleep in zebrafish models of Usher syndrome type 2a

**DOI:** 10.64898/2026.03.09.710525

**Authors:** Jessie M. Hendricks, Vikash Choudhary, Charles R. Heller, Mel van Gemert, Daniël L.A.H. Hornikx, Sanne Broekman, Theo Peters, Jan Zethof, Jonathan Hensman, Camiel J.F. Boon, Drew N. Robson, Jennifer M. Li, Rianne Ligterink, Annemieke Kleinhout-van Vuuren, Silvia C. Endenburg, H. Myrthe Boss, Rob W.J. Collin, Juriaan R. Metz, Erik de Vrieze, Erwin van Wijk

**Affiliations:** Department of Otorhinolaryngology, Radboud University Medical Center, Nijmegen, The Netherlands; Donders Institute for Brain, Cognition and Behaviour, Radboud University Nijmegen, Nijmegen, The Netherlands; Max Planck Institute for Biological Cybernetics, Tübingen, Germany; Department of Human Genetics, Radboud University Medical Center, Nijmegen, The Netherlands; Department of Biomolecular Chemistry, Institute for Molecules and Materials, Radboud University Nijmegen, Nijmegen, The Netherlands; Department of Plant & Animal Biology, Radboud Institute for Biological and Environmental Sciences, Radboud University Nijmegen, Nijmegen, The Netherlands; Department of Ophthalmology, Amsterdam University Medical Centers, University of Amsterdam, Amsterdam, The Netherlands; Department of Ophthalmology, Leiden University Medical Center, Leiden, The Netherlands; Department of Neurology, Hospital Gelderse Vallei, Ede, The Netherlands

## Abstract

Usher syndrome type 2A (USH2a), the most common form of hereditary deaf-blindness, is frequently accompanied by fatigue and poor sleep quality. As these sleep problems occur independently of visual decline, it is hypothesized that the *USH2A*-encoded protein usherin regulates sleep and circadian rhythmicity via an extra-retinal mechanism. *Ush2a* knockout zebrafish models were utilized to investigate this hypothesis. Immunohistochemical analysis demonstrated usherin localisation in pineal gland photoreceptor cells in wild-type larvae, alongside the USH2 complex proteins Adgrv1 and Whrna. Cross-species transcriptomic and proteomic analyses confirmed *USH2A* expression in all mammalian pineal gland tissues studied. Circadian clock gene expression was measured over 24 h and showed preserved oscillatory patterns in wild-type and mutant zebrafish. *Ex vivo* superfusion of pineal glands revealed sustained circadian melatonin release with comparable phase and period to controls, although potential differences in absolute melatonin levels could not be excluded. Despite intact clock gene expression and melatonin release in *ush2a* mutants, behavioural classification over 24-h recordings revealed altered sleep-wake behaviour: *ush2a* mutants displayed elevated daytime sleep and significantly prolonged and more variable sleep latency. The dissociation between intact molecular rhythms and abnormal sleep behaviour likely implicates that usherin plays a role in sleep-wake regulation independent of the circadian pacemaker and melatonin synthesis. These findings suggest a novel role of usherin in the pineal gland and establish a mechanistic link between usherin dysfunction and sleep disturbances, providing a biological basis for the fatigue and sleep problems reported in USH2a patients.

## INTRODUCTION

Usher syndrome (USH) is the most common form of hereditary deaf-blindness, accounting for about 50-70% of all inherited cases (Bahena et al., 2022; Karuntu et al., 2025; Machado et al., 2025; Slijkerman et al., 2017). It affects approximately 400,000 individuals worldwide. Pathogenic variants in at least 11 different genes have been identified to cause USH (Riazuddin et al., 2025; Ullah et al., 2025). Due to its genetic and clinical heterogeneity, USH is classified into four types: USH1-4. Approximately two-thirds of Usher patients suffer from USH2, which is characterized by moderate to severe congenital hearing impairment and the onset of progressive vision loss due to retinitis pigmentosa (RP) around the second decade of life. Up to 85% of all USH2 patients are genetically classified as Usher syndrome type 2a (USH2a), caused by bi-allelic pathogenic variants in the *USH2A* gene (Millán et al., 2010; Yan & Liu, 2010). Besides Usher syndrome, pathogenic variants in the *USH2A* gene are also the most common cause of non-syndromic retinitis pigmentosa (Karuntu et al., 2025; Verbakel et al., 2018). The *USH2A* gene encodes usherin, a large transmembrane protein. In the cochlea, it is a component of the ankle link complex of the auditory hair bundles and it is furthermore located at the synapse of the auditory hair cells and in the spiral ganglion neurons (van Wijk et al., 2006). In the retina, usherin is located at the photoreceptor periciliary membrane, bridging the gap between the apical inner segment and the connecting cilium. Usherin is known to be essential for cochlear hair cell development and retinal photoreceptor maintenance (Liu et al., 2007; Mathur & Yang, 2015).

Beyond the congenital hearing impairment and progressive loss of visual function, several other, mostly subclinical, comorbidities have been reported in individuals with USH2a. These co-morbidities include olfactory defects (Jansen et al., 2016; Ramos et al., 2019; Zrada et al., 1996), reduced touch sensitivity (Frenzel et al., 2012; Schwaller et al., 2021), and fatigue (Ehn et al., 2020; Wahlqvist et al., 2013). Fatigue, a prevalent and debilitating symptom in individuals with USH2a, has historically been attributed to the strain of coping with a dual sensory loss or the associated mental health challenges. However, a questionnaire-based sleep study revealed that USH2a patients not only experience fatigue but also suffer from significantly poorer sleep quality, a higher incidence of sleep disorders, and elevated daytime sleepiness compared to unaffected controls (Hendricks et al., 2023). A follow-up study utilizing actigraphy and a sleep diary confirmed that USH2a patients suffer from a significantly longer sleep latency, reduced sleep and rest quality, and greater day-to-day variability in sleep efficiency and sleep latency (Hendricks et al., 2024). Together, these results could give rise to the assumption that impaired sleep in USH2a potentially implicates disturbances in light-dependent circadian regulation.

Light is the principal zeitgeber for circadian rhythms, synchronizing the internal clock with the external environment (LeGates et al., 2014; Takahashi & Zatz, 1982). In humans, this process is primarily mediated by intrinsically photosensitive retinal ganglion cells (ipRGCs), which express the photopigment melanopsin and transmit light information directly to the suprachiasmatic nucleus in the brain (Berson et al., 2002). Therefore, circadian dysfunctions are more prevalent in individuals with complete blindness, where the lack of retinal light input leads to free-running rhythms and sleep–wake disorders (Hartley et al., 2018). Based on this knowledge, one might hypothesize that the sleep problems in USH2a patients have a similar underlying cause. However, it is crucial to note that although RP may eventually lead to legal blindness, individuals with USH2a typically preserve functional central vision well into the second to fourth decade of life, and even in late stages many individuals retain some degree of residual light perception (Pierrache et al., 2016). Importantly, the study of Esquiva et al. (2013) showed that ipRGCs remain largely preserved at advanced stages of non-syndromic RP, even after photoreceptor degeneration, suggesting that circadian photoreception may remain functional. And consistent with these reports, both the questionnaire and the actigraphy-based USH2a sleep studies found no correlation between disease progression and the sleep problems (Hendricks et al., 2024; Hendricks et al., 2023). Hence, these findings indicate that the sleep disturbances are intrinsic to USH2a, occurring independently of the severity of the visual impairment.

To study the function of USH-associated proteins in the retina and cochlea, the most commonly used animal models are zebrafish and mice. Knock-out and knock-in models are available for nearly all genes associated with USH, providing critical insights into disease mechanisms and enabling the evaluation of therapeutic interventions under development. While USH mouse models are particularly valuable for investigating cochlear phenotypes (Stemerdink et al., 2021), they fail to fully recapitulate the human retinal degeneration phenotype, likely due to anatomical differences in photoreceptors (Slijkerman et al., 2015). Therefore, most insights into mechanisms underlying Usher syndrome-associated retinal disease have been obtained from studies using zebrafish models. Zebrafish *ush2a* mutants display early-onset retinal dysfunction and present with impaired intracellular transport of photopigments (Dona et al., 2018; Reurink et al., 2022). Despite the extensive use of USH models for cochlear and retinal research and therapy development, sleep and circadian phenotypes associated with USH have not been investigated in any animal model to date. Importantly, zebrafish are exceptionally well suited to address this gap: unlike rodents, they are diurnal, displaying robust circadian and sleep-like behaviours from early larval stages. Additionally, the expression of circadian clock genes is highly conserved between zebrafish and humans (Livne et al., 2016; Rihel et al., 2010; Vatine et al., 2011). Together, these features position zebrafish as an excellent model for exploring the link between *USH2A* dysfunction, circadian regulation, and sleep–wake behaviour.

Zebrafish do not rely on the suprachiasmatic nucleus (SCN) as the central circadian pacemaker and this unique organization makes zebrafish particularly suitable for dissecting the direct effect of light perception and photoreceptor dysfunction on circadian regulation and sleep-wake behaviour. In mammals, environmental light is detected by the retina and transmitted via specialized retinal ganglion cells to the SCN, where it entrains the molecular clock that regulates systemic rhythms (Berson et al., 2002). In contrast, zebrafish possess a more decentralized system in which multiple tissues, including the retina, liver, and heart, contain their own directly light-entrainable clocks (Cahill, 1996; Frøland Steindal & Whitmore, 2019). At the core of the zebrafish circadian system is the pineal gland, a small neuroendocrine organ located at the dorsal edge of the diencephalon (Toyama et al., 2009). The zebrafish pineal gland is directly photosensitive and produces melatonin in a circadian manner (Appelbaum et al., 2009; Cahill, 1996; Ziv et al., 2007). Its main cell type is the pineal photoreceptor cell, also named pinealocyte. Zebrafish pinealocytes share key structural and molecular features with retinal photoreceptors, including the presence of outer segments with stacked lamellae and expression of rhodopsin (Falcón, 1999; Laurà et al., 2012). Studies indicate that pineal and retinal photoreceptors share a common ancestral origin, having evolved from a single primordial photodetector cell (Ekström & Meissl, 2003; Klein, 2006; Staudt et al., 2019).

Given the shared evolutionary and molecular features of pineal and retinal photoreceptors, together with the presence of usherin in retinal photoreceptor cells, we hypothesized that usherin contributes to the regulation of circadian rhythmicity and sleep-wake behaviour. Here, we show that usherin localises to the pineal gland and that *ush2a* knockout zebrafish display increased daytime sleep and prolonged sleep latency despite only mild visual dysfunction. These findings support a direct role for usherin in sleep regulation and suggest a potential extra-retinal mechanism underlying USH2a-associated sleep problems.

## METHODS

### Fish maintenance and husbandry

All zebrafish were raised and maintained according to the guidelines (Aleström et al., 2019; Westerfield, 2007). Adult zebrafish were maintained at 28,5°C on a 14h:10h light/dark (LD) cycle, with lights (400 lux white LED) on at 9 AM and lights off at 11 PM. Embryos were obtained by natural spawning after the onset of light and maintained in 10cm Petri dishes in E3 medium which was refreshed every day and incubated in the same temperature and light regimen as the adult zebrafish. In this study, two previously published *ush2a* zebrafish models were utilized. *ush2a^b1245^*, an AB strain zebrafish carrying a frameshift mutation in *ush2a* exon 71 (c.15520_1552delinsTG; p.Ala5174fsTer), and *ush2a^rmc1^*, a Tüpfel long fin (TL) strain zebrafish carrying frameshift mutation in *ush2a* exon 13 (c.2337_2342delinsAC; p.Cys780GlnfsTer32) (Dona et al., 2018). Heterozygous fish were incrossed, raised, and genotyped, resulting in F1 homozygous mutants and wild-type siblings. Both the homozygous mutants and wild-type siblings were incrossed to generate F2 offspring for the larval experiments.

### Immunohistochemistry on zebrafish larvae

Immunohistochemistry was performed on 5 dpf larval zebrafish of the following genotypes: *ush2a^b1245^*, *ush2a^rmc1^*, and age and strain-matched wild-types. Euthanized larvae were rinsed in 10% sucrose, embedded in OCT compound (Tissue-Tek, Sakura) and frozen in melting isopentane. 7µm sagittal cryosections were made. After drying of the cryosections, a pre-fixation or post-fixation immunohistochemical procedure was followed. Pre-fixation procedure: sections were washed with PBS and subsequently fixed with 4% PFA for 10 minutes. Afterwards, the sections were washed three times with PBS, followed by 20 minutes of permeabilization in 0.1% Tween-20 in PBS. Then, sections were blocked for one hour with 10% Normal Goat serum (NGS) and 10% Bovine Serum Albumin (BSA) in PBS. Primary antibodies were incubated overnight at 4°C in blocking solution. The following primary antibodies were used: rabbit anti-usherin (C-terminal, 1:1000, DZ01481, Boster Bio) and mouse anti-rhodopsin (1:5000, NBP2-59690, Novus Biologicals) or mouse anti-centrin (1:500, 04-1624, Sigma-Aldrich). After six rounds of washing with PBS, the secondary antibodies (1:800; goat anti-mouse Alexa Fluor 488, A11029, Invitrogen; goat anti-rabbit Alexa Fluor 568, A11011, Molecular Probes) and DAPI (1:8000) were incubated in blocking buffer for 2 hours at room temperature (RT). Sections were washed 6 times with PBS before mounting with Prolong Gold Anti-fade (P36930, Thermo Fisher Scientific). Post-fixation procedure: unfixed cryosections were washed with PBS followed by 20 minutes of permeabilization in 0.01% Tween-20 in PBS. Then, sections were blocked with 10% NGS and 10% BSA in PBS. Primary antibodies were incubated overnight at 4°C in blocking solution. The following primary antibodies were used with the post-fixation procedure: guinea pig anti-usherin (N-terminal, 1:500, kindly provided by the Westerfield lab, University of Oregon), mouse anti-centrin (1:500, 04-1624, Sigma-Aldrich), mouse anti-rhodopsin (1:5000, NBP2-59690, Novus Biologicals), rabbit anti-Adgrv1 (1:100, DZ41032, Boster Bio) or rabbit anti-Whrna (1:500, 42700002, Novus Biologicals). After six rounds of washing with PBS, the secondary antibodies (1:800; goat anti-mouse Alexa Fluor 488, A11029, Invitrogen; goat anti-rabbit Alexa Fluor 568, A11011, Molecular Probes; donkey anti-guinea pig Alexa Fluor 568, A11057, Molecular Probes) and DAPI (1:8000) were incubated in blocking buffer for 2 hours at RT. Sections were washed 6 times with PBS before fixing with 4% PFA for 10 minutes. After washing 3 times with PBS, the sections were mounted with Prolong Gold Anti-fade (P36930, Thermo Fisher Scientific). Images were taken using a Zeiss Axio Imager fluorescence microscope equipped with an AxioCam MRC5 camera (Zeiss).

### *USH2A* gene expression in higher vertebrates

To study *USH2A* gene expression in retina and pineal gland of higher vertebrates, the SNE-NICHD Transcriptome Profiling web page and the Non-Human Primate Cell Atlas (NHPCA) were utilized (Chang et al., 2020; Han et al., 2022). From the SNE-NICHD database, the following *USH2A* gene expression data were extracted: zebrafish (*Danio rerio*), mouse (*Mus musculus*), rat (*Rattus norvegicus*), and rhesus macaque (*Macaca mulatta*). For these four species, pineal gland, retina, and mixed tissue control samples were included, collected during both day and night. For human (*Homo sapiens*), pineal gland day and night samples were available and included. From the NHPCA, *USH2A* single-cell gene expression data from 45 tissues of cynomolgus macaque (*Macaca fascicularis*) were visualized in a UMAP plot via the gene expression tool on their website. To visualize the evolutionary distance between the abovementioned species and pig (*Sus scrofa*, see next section), a phylogenetic tree was constructed as follows: *USH2A* amino acid sequences were retrieved from UniProt (The UniProt Consortium, 2024). Multiple sequence alignment was performed with MAFFT version 7 using default FFT-NS-i settings (Katoh et al., 2017). The phylogenetic tree was generated using the default Neighbour-Joining method and JTT substitution model, exported as Newick file and imported in Interactive Tree Of Life (iTOL) version 7.2 for visualization (Letunic & Bork, 2024).

### Western blot on porcine retina and pineal gland

Whole pig eyes were received from the local abattoir, directly placed on ice after the animals were sacrificed and dissected within 4 hours. An incision was made with a scalpel and the sclera was cut with scissors. The lens and vitreous humor were removed and the retina was separated from the optic nerve. The retina was snap frozen in liquid nitrogen and stored at - 80°C. Porcine pineal glands were dissected at the PIGMOD Center in Liběchov, Czech Republic. Pineal glands from a 7-month-old female pig sampled six hours after light-on, and a 4.5-month-old male pig sampled 8 hours after light-on, were snap frozen in liquid nitrogen and stored at −80°C. Tissue lysis was performed as follows: Tissues were thawed, cut into small pieces with a scalpel and washed with ice cold PBS. After removing the PBS, 1ml cell lysis buffer (150mM NaCl, 50mM Tris-HCl, 5mM EDTA, 1% triton, 0.1% SDS (15553027, Invitrogen), Protease inhibitor cocktail (11697498001, Roche) in MQ) was added to a sample containing a whole retina, and 200µl cell lysis buffer was added to half a pineal gland. The tissue in cell lysis buffer was incubated on ice for 1 hour while shaking. Then, the samples were centrifuged for 5 minutes on 400xg at 4°C. The supernatant was transferred to a new Eppendorf and centrifuged at 15,000 rpm for 15 minutes at 4°C. The supernatant was designated the *soluble fraction*. The *insoluble fraction (*pellet) was further extracted by resuspension in 100-200µl cell lysis buffer with 1% additional SDS.

Total protein concentration of all samples was determined with the Pierce BCA Protein Assay Kit (23225, Thermo Scientific). A total of 20µg of protein per sample was mixed with NuPAGE LDS sample buffer (NP0007, Invitrogen) and 100mM DTT. These samples were heated for 10 minutes at 70°C, centrifuged shortly and cooled on ice. Samples and the HiMark Pre-stained Protein Standard (LC5699, Invitrogen) were loaded on a 5% handcasted tris-acetate gel. After running for 3 hours on 60V the proteins were transferred to a PVDF membrane (GE10600023, Merck) using the Mini Blot Module (B1000, Invitrogen), for 2 hours at 325mA constant amperage at 4°C. Then, the membrane was rinsed with PBS, and blocked with 5% BSA in PBS for 1 hour at RT. The primary antibody was custom-made mouse monoclonal antibody generated against the intracellular amino acids EAIMGHNSGLYVDEEDLMN of human usherin. Specificity of this antibody for usherin was validated in a previous publication (Sanjurjo-Soriano et al., 2023). A Basic Local Alignment Search Tool (BLAST) analysis indicated 79% amino acid conservation of this region between human and porcine usherin. The anti-usherin antibody was diluted 1:1000 in 1% BSA in PBS and incubated overnight at 4°C. The next day, the membrane was washed three times for 10 minutes with 0.2% Tween-20 in PBS. The secondary antibody Goat anti-Mouse Alexa Fluor 680 (A-21057, Thermo Fisher Scientific) was diluted 1:20.000 in 1% BSA in PBS and incubated for 1 hour at RT. Then, three rounds of washing were performed, followed by rinsing and storing in PBS. The membrane was imaged with the Odyssey DLx Imager (LICORbio). To visualize the total amount of protein, a Ponceau S staining was performed afterwards. A 0.1% Ponceau S in 5% acetic acid solution was made using Ponceau S powder (P3504, Sigma-Aldrich). The membrane was washed in MQ three times for 1 minute with gentle agitation, followed by immersion of the membrane in the Ponceau S solution for 5 minutes with gentle agitation. Then, the membrane was washed three times in MQ and scanned with the Epson Perfection V330 Photo scanner.

### Gene expression analysis

Four dpf *ush2a^b1245^*, *ush2a^rmc1^*, AB and TL wild-type zebrafish larvae were snap frozen in liquid nitrogen at zeitgeber time (ZT) 0, 4, 8, 12, 16, 20 and 24. Zeitgeber time refers to time relative to the entraining light-dark cycle (ZT0 = lights on; ZT14 = lights off). For each ZT, 4 pools of 3 larvae were collected per genotype. Samples were stored at −80°C until RNA isolation. Total RNA was isolated using the following procedure: First, the tissue was homogenized with 400μl Trizol reagent (Invitrogen, Carlsbad, USA) and a plastic grinding ball in a Grinding Mill (Retsch GmbH, Germany) for 30s at 20 Hz. After homogenization, samples were kept at RT for five minutes. Next, 80 μl chloroform was added and the solution was mixed by shaking for 15s. Samples were kept at RT for 2 minutes and then centrifuged at 14,000 rpm for 10 minutes at 4°C. The aqueous phase of the samples was transferred to a new Eppendorf. 200 μl isopropanol was added to the aqueous phase and this solution was mixed by inversion of the tube. The solution was stored at −20°C for 1h. Afterwards, the samples were centrifuged for 15 minutes at 14,000 rpm at 4°C. The supernatant was removed, and the pellet washed with 500 μl 75% ethanol and centrifuged 10 minutes at 14,000 rpm at 4°C. After discarding the supernatant, the pellet was air-dried for 10 minutes at RT and then dissolved in 100μl DEPC-treated water. 10 μl 3M NaOAc pH 5.4 and 250 μl 100% EtOH were added to the solution. After mixing by inverting the tube, samples were stored for 1 h at −20°C. Next, the samples were centrifuged for 15 minutes at 14,000 rpm at 4°C, the supernatant was removed, and the pellet washed as described earlier. Finally, the RNA was dissolved in 15 μl DEPC-treated water. The concentration and quality of RNA were measured using a Nanodrop spectrometer at 260 and 280 nm wavelength (Nanodrop, Wilmington, DE, USA). After DNase treatment (18068015, Invitrogen), cDNA was synthesized: 1μl random primers (250 ng/μl), 1μl 10 mM dNTP mix, 4μl 5×1st strand buffer, 1 μl 0.1 M DTT, 1μl RNase Inhibitor (10 U/μl), 0.5μl Superscript II (200 U/μl) (Invitrogen, Carlsbad, USA) and 0.5 μl DEPC-treated H_2_O were added to the samples. The resulting mix was incubated for 10 minutes at 25°C for primer annealing and then 50 minutes at 42°C for first strand synthesis. After this, enzymes were inactivated by incubating samples at 70°C for 15 minutes. Finally, H_2_O was added to dilute the samples five times for the qPCR reaction. The gene expression of the following clock genes was measured by a quantitative PCR (qPCR): *aanat2*, *bmal1a*, *clock1a*, *cry1a*, *cry2a* and *per2*. *βactin* and *rpl13* were used as housekeeping genes (Casadei et al., 2011; McCurley & Callard, 2008). All primer sequences are listed in supplementary table s1. For each qPCR reaction, 16μl PCR mix (containing 2x SYBR green mix (BioRad, Hercules, USA), 10µM forward and reverse primers, and MQ) was added to 4 μl of diluted cDNA. The qPCR reaction (3 min 95°C, 40 cycles of 15s 95°C and 1 min 60°C) was carried out using a CFX 96 qPCR machine (BioRad, Hercules, USA). qPCR data were analysed in Bio-Rad CFX Manager. Expression was normalized for *βactin* and *rpl13*. Routine quality check procedures were followed: these include melting curve analyses, no-RT and no-template controls (Vandesompele et al., 2002). Gaussian distribution was assumed, and an unpaired t-test was performed for every timepoint, along with a correction for multiple comparisons using the Holm-Šídák method, to compare the relative gene expression of mutant and control samples.

### Pineal gland superfusion and melatonin measurements

To investigate melatonin release, two-year-old *ush2a^b1245^*mutant zebrafish and wild-type siblings were dissected for *ex vivo* superfusion of the pineal gland. Fish were anesthetized in 0.1% (v/v) 2-phenoxyethanol. The skullcap was surgically removed under a dissection microscope and the brain was extracted. The brain was further dissected using a conservative approach to preserve the integrity of the target region. The dissected tissues were temporarily stored in a well-plate with superfusion medium in normal light conditions until the pineal glands of all fish were successfully dissected. Then, the dissected tissues were placed on a filter in 15µl superfusion chambers around ZT9, starting the superfusion of the tissues with the medium. At ZT14, the normal light offset time, the light was turned off and sample collection (the outflowing solution, or superfusate) was started. The light was kept off for the whole further duration of the experiment.

The culture medium used was modified from previously described formulations of balanced salts and amino acids (Besharse & Dunis, 1983; Cahill & Besharse, 1991; Cahill, 1996). It consisted of 106mM NaCl, 2mM KCl, 1.8mM CaCl_2_, 1mM MgCl_2_, 35mM NaHCO_3_, 1mM NaH_2_PO_4_, 5mM D(+)glucose, 0.1mM L(+)ascorbic acid, 1.1mM 5-hydroxy-L-tryptophan (H9772, Sigma-Aldrich), Penicillin-Streptomycin (P0781, Sigma-Aldrich), and 14 amino acids (Supplementary table S2). The medium was prepared in the following order to prevent precipitation: KCl, NaH_2_PO_4_, NaHCO_3_, L(+)ascorbic acid and D(+)glucose were dissolved in water. While mixing, 5-hydroxy-L-tryptophan and Penicillin-Streptomycin were added from 100x stock solutions. After the medium was saturated for 1 hour with carbogen (95% O_2_/5% CO_2_), all amino acid stocks were added, followed by CaCl_2_ and MgCl_2_ stocks. The medium had a pH of 7.4 and an osmolality of 290 mOsmol/kg, closely matching physiological conditions. The medium was stirred and saturated with carbogen during the experiment.

Culture medium was delivered with a flow rate of 270µl/h with a multichannel peristaltic pump (Watson–Marlow, Falmouth, Cornwall, UK). Medium and tissues were maintained at RT throughout the experiment. Superfusate samples from each chamber were collected in 2-hour intervals. Samples were collected in the dark during the experiment after a maximum of 16 hours at RT and stored at −20°C until melatonin measurements. 2.5 days after the start of the experiment, the tissues were collected from the chamber to check the viability with the CellTiter-Glo 3D Cell Viability Assay (G9681, Promega) according to the standard protocol. No differences were found between the viability of *ush2a^b1245^* and control pineal glands.

Melatonin concentrations in the superfusates were measured using liquid chromatography-tandem mass spectrometry (LC-MS/MS) (van Faassen et al., 2017). Concentrations (pg/ml) were converted to secretion rates (pg/hour) based on the flow rate. To assess statistical differences at individual time points, unpaired t-tests were performed, with multiple comparisons corrected using a false discovery rate approach. For each sample, melatonin concentrations were normalized to the mean across all time points to obtain relative melatonin levels. The relative melatonin concentrations were used to fit a sine wave function with a nonzero baseline: Y = Amplitude * sin((2π * X / Wavelength) + Phaseshift) + Baseline. Fitting was performed separately for control and mutants and without constraints. The resulting best-fit parameters amplitude, wavelength, phaseshift and baseline were compared between the two genotypes using a comparison-of-fits analysis. All melatonin visualizations and statistical analyses were performed in GraphPad Prism 10.4.1 for Windows (GraphPad Software, Boston, Massachusetts USA, www.graphpad.com)

### Behavioural analysis

The circadian sleep-wake behaviour of *ush2a^b1245^* mutant larvae and control offspring (derived from incross of wild-type siblings) was assessed using the DASHER behavioural acquisition system (Dynamic Addressable System for High-speed and Enhanced Resolution) (Choudhary et al., 2023). Behavioural experiments were conducted in larvae with an AB background, as this strain is widely used in sleep research and exhibits lower anxiety-induced behaviour (Egan et al., 2009), faster habituation, and more pronounced light-dark responses compared to other strains (van den Bos et al., 2017). Embryos were raised in the standard LD cycle. Behavioural recordings began at 5 dpf using two DASHER setups in parallel: one capable of recording up to 16 larvae simultaneously, and the other up to 20. Individual larvae were placed in circular chambers (28 mm diameter, 1 mm height) filled with fresh E3 medium. Chambers were sealed with glass coverslips to prevent evaporation. Half of the chambers were loaded with *ush2a^b1245^*mutant larvae, and the other half with controls. Larvae were recorded in the DASHER setup for 24 hours in normal LD conditions. After the first recording session, the chambers were emptied and reloaded with the remaining larvae from the same clutch, which were measured at 6 dpf. A second, independent round of data collection was performed as a biological replicate, using embryos from a separate clutch. After each experiment, the larvae were checked on viability and morphological abnormalities. For detailed methods on data acquisition, eye tracking, and sleep-state determination, see Choudhary et al. (2023).

Locomotor and eye tracking were successfully completed for 70 *ush2a^b1245^* mutant larvae and 66 controls. Each individual larva’s behavioural state profile was visualized, along with a density plot representing the group’s state probabilities. For statistical analysis, the cumulative durations of each behavioural state were calculated per larva for the light period (ZT0-14, 9:00AM-11:00PM) and the dark period (ZT14-24, 11:00PM-9:00AM), and compared between genotypes using the Wilcoxon rank-sum test. Sleep latency was also calculated for each larva, defined as the time taken to transition from wakefulness to any sleep substate after the light offset at ZT14. Sleep latencies were compared between genotypes using the Wilcoxon rank-sum test, and variability in sleep latency was assessed with Levene’s test. All DASHER output data visualizations and statistical analyses were performed in Python 3.12 within the Spyder 5.5.1 integrated development environment, part of the Anaconda distribution (Python Software Foundation, https://www.python.org/; Spyder IDE, https://www.spyder-ide.org/; Anaconda Inc., https://www.anaconda.com/).

## RESULTS

### Usherin is expressed in the photosensitive pineal gland of wild-type zebrafish

Following the evolutionary relationship between the zebrafish retinal photoreceptors and pinealocytes, we first investigated if the *ush2a*-encoded protein usherin was also present in the zebrafish pineal gland. Whole 5 dpf zebrafish larvae were serially sectioned and stained, with a particular focus on the brain. In figure 1A, a mid-sagittal hematoxylin and eosin (H&E) stained section from the Zebrafish BioAtlas is shown (Bio-Atlas, 2013; Copper et al., 2018). The pineal gland is located on the dorsal side of the brain, situated between the telencephalon and mesencephalon. In figure 1B, a corresponding mid-sagittal section is shown, stained for the nuclear marker DAPI (blue), the pineal gland marker rhodopsin (green), and the C-terminal domain of usherin (red). A distinct usherin signal was observed in the pineal gland region. Higher magnification images of the pineal gland of AB wild-type and *ush2a^b1245^* larvae are shown in figure 1C. Usherin immunoreactivity was clearly detected in pineal gland of wild-type larvae but was not detectable in *ush2a ^b1245^*mutants using the C-terminal antibody.

**Figure 1.**
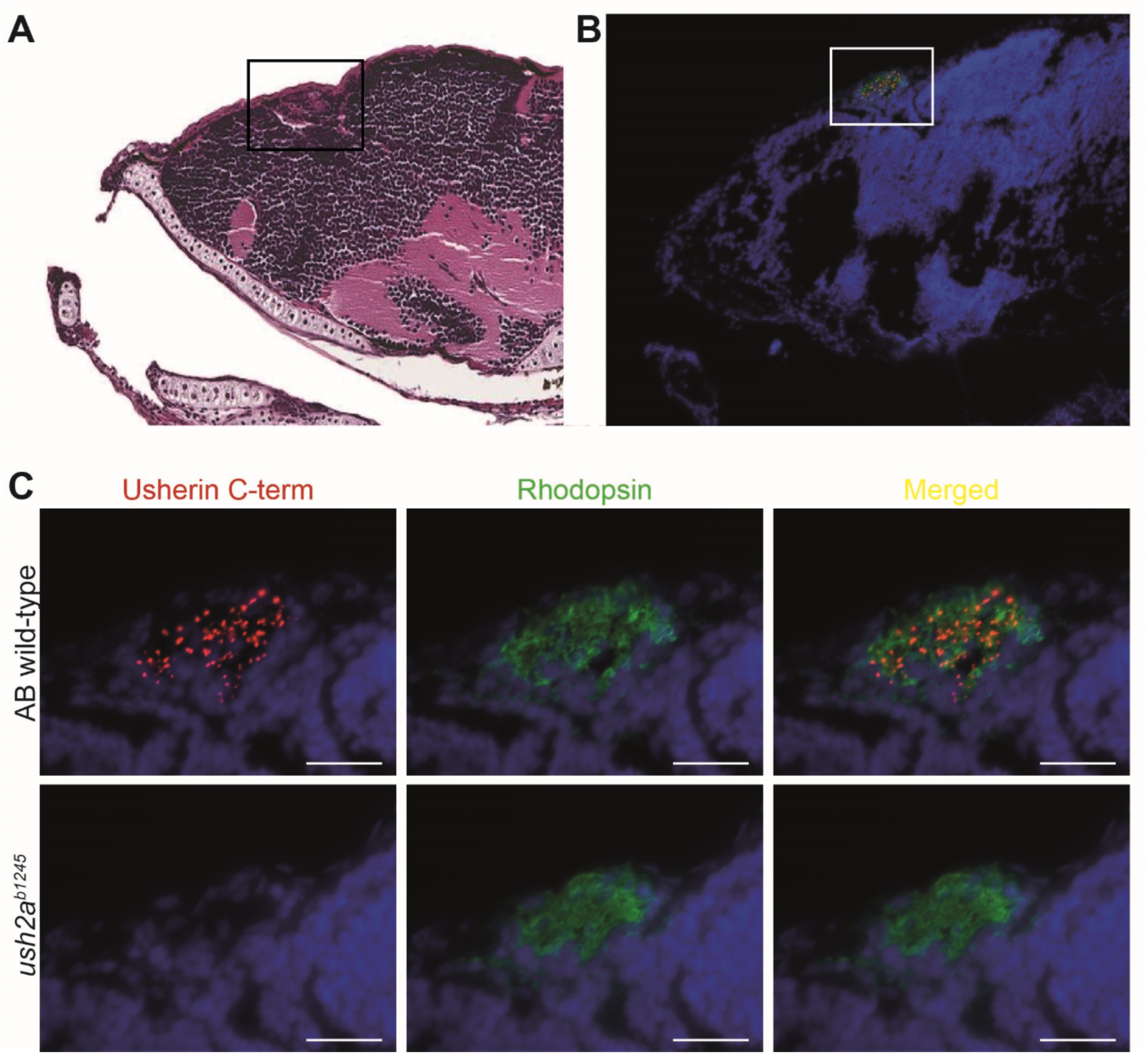
Usherin is expressed in the pineal gland of wild-type zebrafish larvae and is absent in *ush2a ^b1245^* mutants. **A)** Mid-sagittal H&E-stained section from the Zebrafish BioAtlas of a 6 dpf larva (Bio-Atlas, 2013; Copper et al., 2018). The pineal gland (indicated by the box) is located on the dorsal side of the brain, situated between the telencephalon and mesencephalon. **B)** Corresponding mid sagittal section from 5 dpf wild-type larva labelled with DAPI (blue), anti-usherin C-terminus (red), and anti-rhodopsin (green). The white box indicates the pineal gland region, which is shown at higher magnification in panel C. **C)** Higher magnification of the pineal gland in AB wild-type and *ush2a^b1245^* mutant larvae. Usherin localises to the pineal gland in wild-type but is absent in *ush2a* mutants. Scale bar: 20 µm.

To further define the subcellular localisation of usherin in the pineal gland, a co-staining was performed with centrin, a marker of the photoreceptor basal body and connecting cilium. In AB wild-type larvae, usherin localised in close proximity to centrin (Figure 2A), which is consistent with its localisation at the photoreceptor connecting cilium in the retina. As expected, usherin C-terminal staining was absent in the *ush2a^b1245^* mutant. Despite the loss of the C-terminal part of the protein, weak N-terminal usherin immunoreactivity was still observed in *ush2a^b1245^* mutants, indicating residual truncated usherin protein is present (Figure 2B). Usherin was also detected in the pineal gland of wild-type larvae of the TL strain, while *ush2a^rmc1^* mutants, which carry a truncating mutation in the extracellular region, lacked both C- and N-terminal usherin staining in the pineal gland (Supplementary Figure S1). These results are consistent with previous observations in retinal photoreceptors cells (Dona et al., 2018).

**Figure 2.**
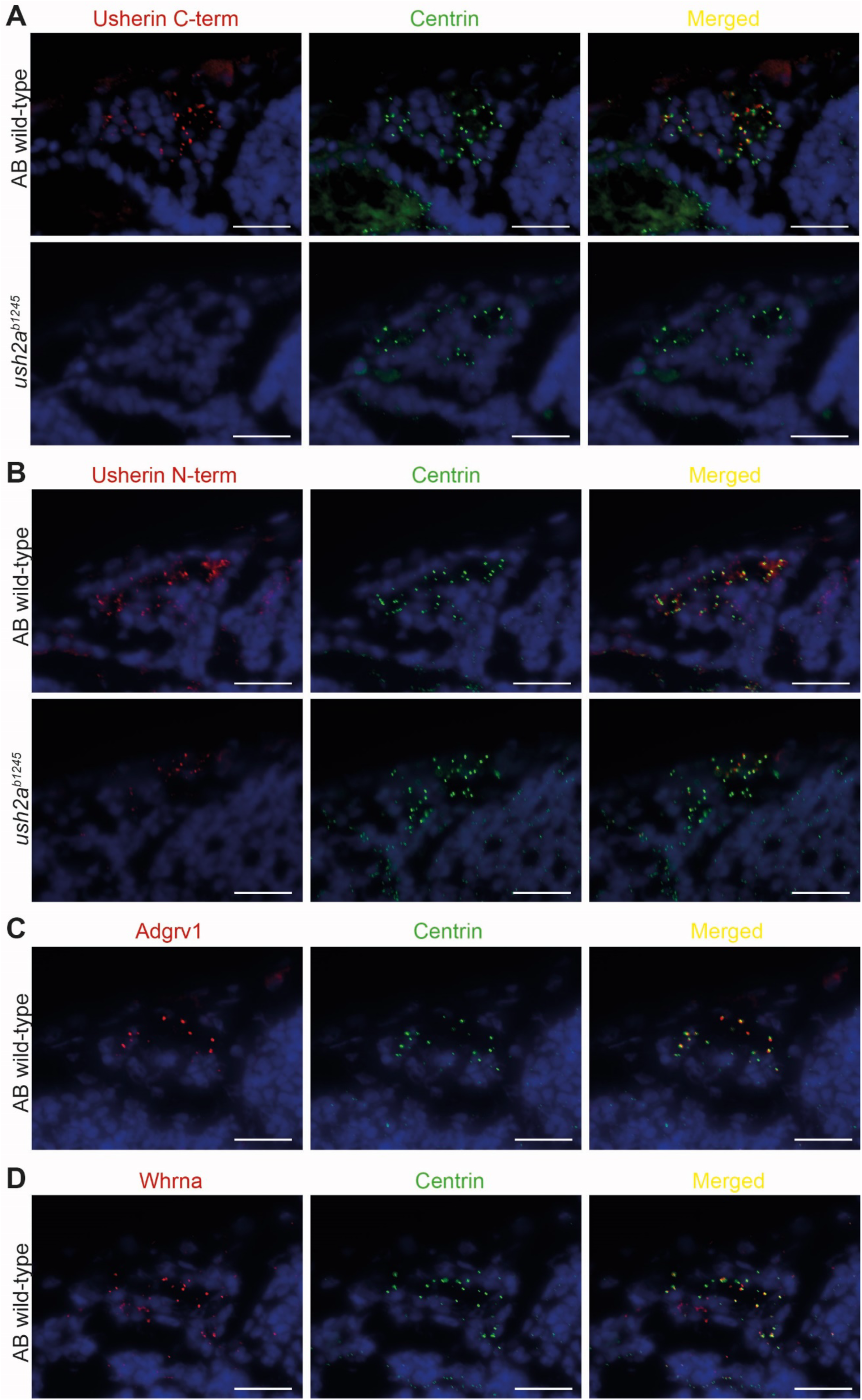
The USH2 complex localises to the connecting cilium of pineal photoreceptor cells. **A)** Immunohistochemical labelling of the pineal gland in AB wild-type and *ush2a^b1245^* larvae showing DAPI (blue), usherin C-terminus (red), and centrin (green). In wild-type larvae, usherin localises in the vicinity of centrin. No usherin C-terminal signal is detected in *ush2a^b1245^* mutants. **B)** Staining for usherin N-terminus (red) in the same genotypes shows reduced but detectable signal in *ush2a^b1245^* mutants **C)** Immunohistochemical labelling of Adgrv1 (red) in the pineal gland of AB wild-type larvae showed similar colocalisation with centrin. **D)** Immunohistochemical labelling of Whrna (red) in the pineal gland in AB wild-type larvae showed similar colocalisation with centrin. Scale bar: 20 µm.

In zebrafish retinal photoreceptors, usherin forms a protein complex with Adgrv1 (associated with Usher syndrome type 2c) and Whrna/b (associated with Usher syndrome type 2d), which is essential for the docking of transport vesicles to transport proteins from the inner to the outer segments of the photoreceptors. Having established that usherin was detected in the vicinity of the cilium and basal body marker centrin, similar to what has been previously observed in retinal photoreceptor cells, the next step was to determine whether the interaction partners of usherin are also present in the pineal gland. Therefore, an immunohistochemical analysis was performed on wild-type larvae using antibodies directed against Adgrv1 and Whrna. Both proteins were detected in the pineal gland, with localisation patterns similar to usherin, indeed confirming that other USH2 complex members are also present (Figure 2C and 2D).

### Usherin is expressed in the mammalian pineal gland

To determine whether *USH2A* expression in the pineal gland is conserved across the vertebrate evolutionary lineage, publicly available transcriptomic data of zebrafish, mouse, rat, non-human primate (*Macaca mulatta*), and human were analysed (Figure 3A). As a reference, *USH2A* expression in the retina and mixed tissue control samples was studied too. As expected, *USH2A* transcripts could be detected in retinae of all species examined, while it was not or minimally expressed in the mixed tissue control samples. Interestingly, *USH2A* was expressed in the pineal gland across all species examined. No consistent day-night patterns in *USH2A* transcript abundance were observed, even when accounting for the diurnal or nocturnal nature of the species (Figure 3B-F). Single-cell transcriptomic data from a non-human primate (*Macaca fascicularis*) confirmed *USH2A* expression in both the pineal gland and retina cell clusters, with no *USH2A* expression detected in the other 43 tissues (Figure 3G). Note that cochlear tissue is not included in this dataset.

**Figure 3.**
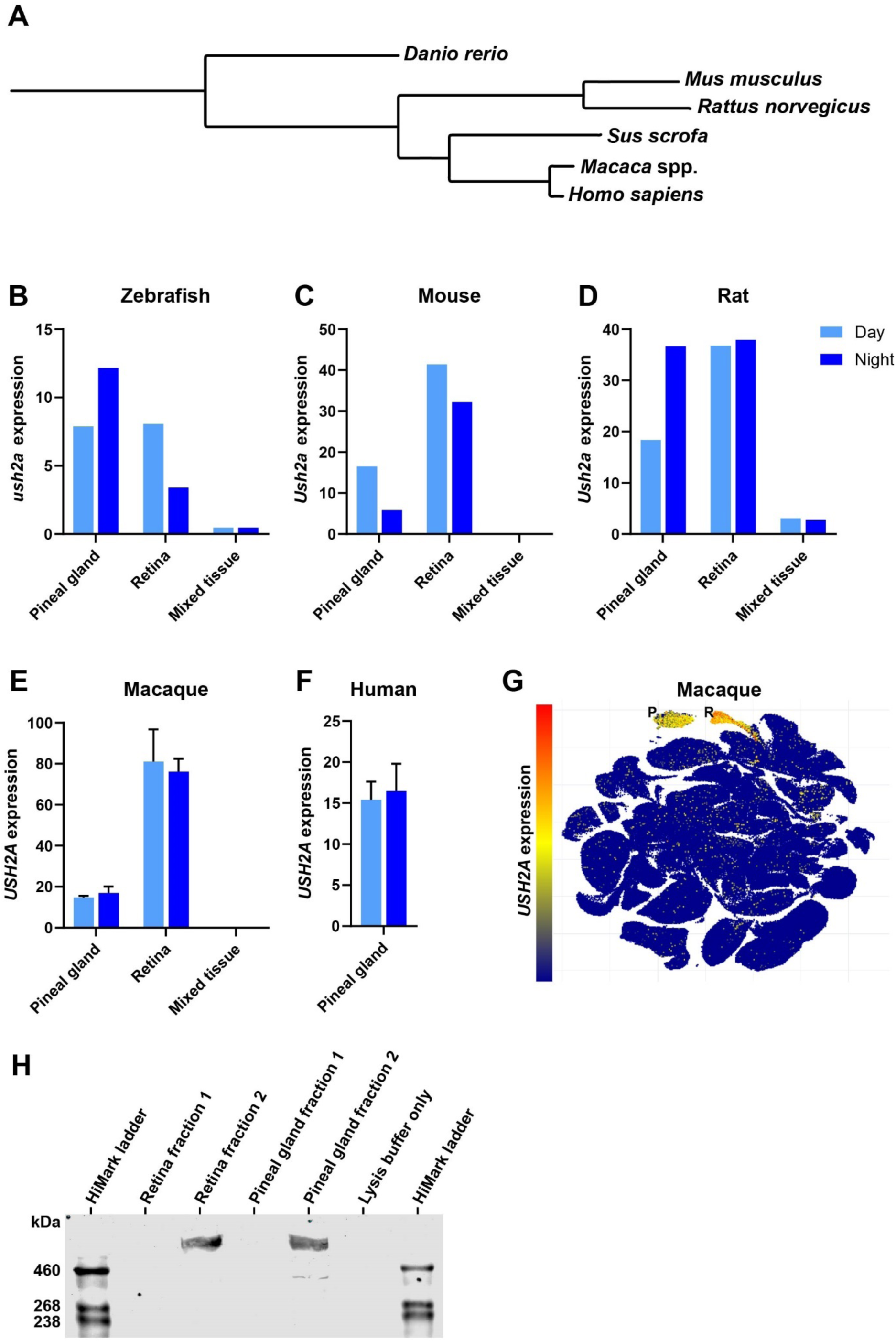
Evolutionarily conserved expression of *USH2A* and usherin in the pineal gland. **A)** Phylogenetic tree showing the evolutionary relationships between species included in the transcriptomic and proteomic analyses. spp. = multiple species (*Macaca mulatta* and *Macaca fascicularis*). Branch lengths are proportional to evolutionary distance. **B–F)** *USH2A* abundance (transcripts per million) in pineal gland, retina, and mixed tissue control samples collected during the day (light blue) and night (dark blue). *USH2A* transcripts are present in both retina and pineal gland across all species examined, with no consistent day–night pattern. **G)** Single-cell transcriptomic map (UMAP) of macaque showing *USH2A* expression restricted to pineal gland (P) and retinal (R) clusters; all other tissues showed no detectable expression. **H)** Western blot detection of usherin protein (∼570 kDa) in the second fraction of porcine retinal and pineal gland lysates. Full blot image and biological replicate are shown in supplementary figure S2.

To verify that the transcriptomic findings translate to protein expression in a large diurnal mammal, western blot analysis was performed on pig tissue lysates. Two fractions were analysed per tissue sample: a soluble fraction, and an insoluble fraction obtained after additional extraction with SDS to release membrane-associated proteins. A high-molecular-weight band of ∼570 kDa was detected in two independent insoluble pineal gland fractions using an anti-usherin antibody, comparable to that observed in retinal lysates (Figure 3H and Supplementary Figure S2). This size is consistent with the predicted molecular weight of full-length usherin (Yu et al., 2020).Together, these findings demonstrate that *USH2A* expression in the pineal gland is conserved from teleosts to mammals, and that usherin protein is present in the mammalian pineal gland despite evolutionary divergence in pinealocyte morphology.

### USH2a zebrafish models have a normal expression of circadian clock genes

To determine whether loss of usherin affects circadian regulation, the expression of circadian clock genes was examined in the two different *ush2a* knockout zebrafish models *ush2a^b1245^*, *ush2a^rmc1^*, and age- and strain-matched wild-type larvae. Expression levels and patterns of *aanat2*, *bmal1a*, *clock1a*, *cry1a*, *cry2a* and *per2* were measured in RNA extracted from whole-larvae collected between 4 and 5 dpf at 4-hour intervals.

In Figure 4, gene expression of *ush2a^b1245^* and AB wild-type zebrafish raised under the conventional 14h:10h light/dark regime are shown. *Aanat2* expression peaked at ZT12 and was lowest between ZT0 and 8. *Bmal1a* and *clock1a* expression also peaked at ZT12, but their expression levels started to increase earlier, starting at ZT4. *Cry1a* and *per2* expression peaked at ZT4, declined during the remainder of the light phase, and reached their lowest levels during the night. While the increase of *cry1a* expression levels started shortly before dawn (ZT0), *per2* expression increased only after light onset. *Cry2a* showed an expression pattern similar to *cry1a* but with a peak shifted toward ZT12. No differences in expression patterns were found in the *ush2a^b1245^* mutants compared to AB wild-type larvae (Figure 4). Clock gene expression analysis of *ush2a^rmc1^* mutants and TL wild-type larvae yielded comparable results (Supplementary Figure S3). Altogether, no difference was observed in the organism-wide expression patterns of circadian clock genes between wild-types and *ush2a* knockout zebrafish larvae.

**Figure 4.**
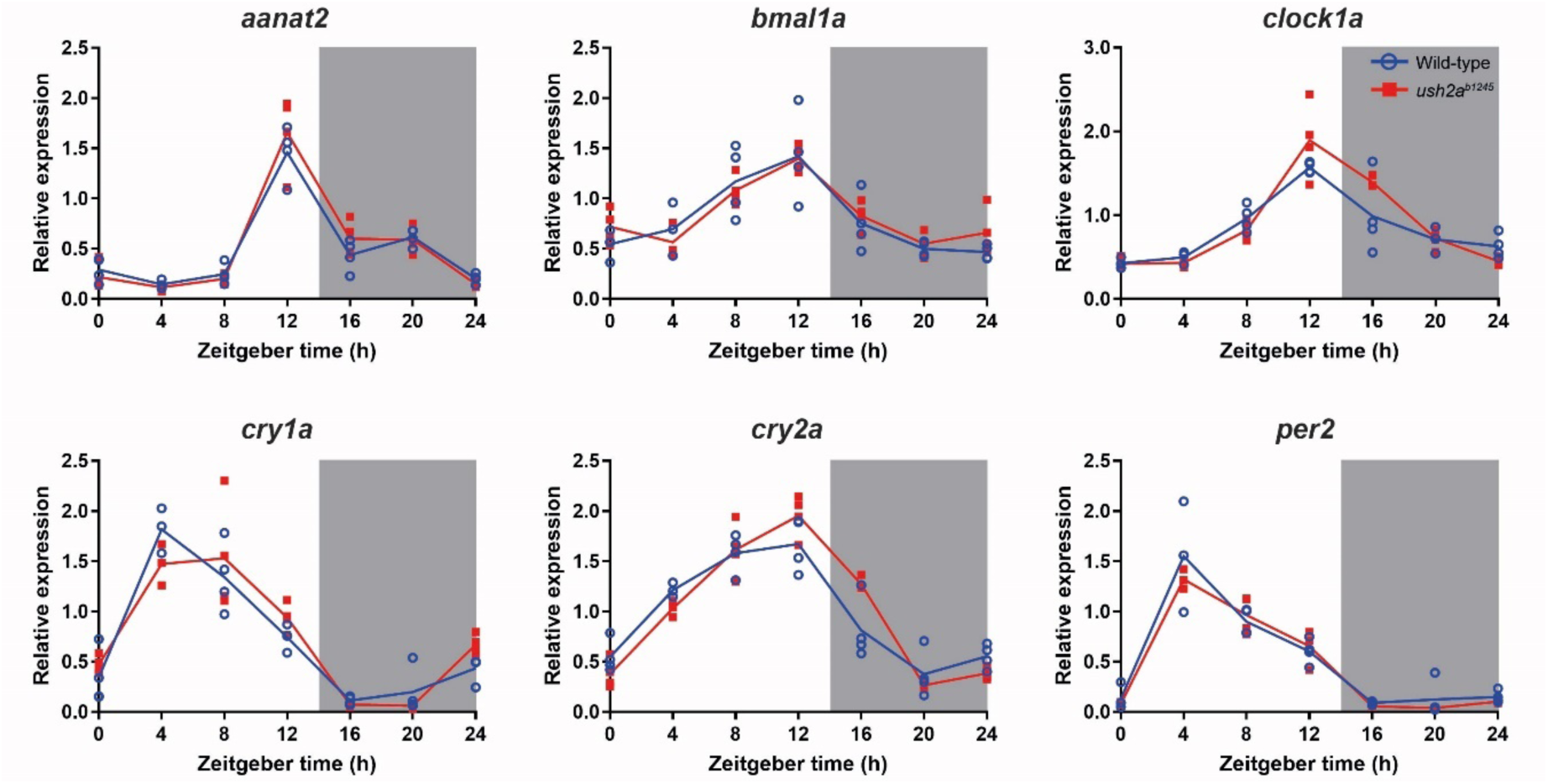
Circadian clock genes expression showed preserved oscillatory patterns in *ush2a^b1245^* larvae. Expression measured over a 24h period at 4-hour intervals, normalized for *βactin* and *rpl13*. Individual measurements are shown as dots (n=4). AB wild-type larvae are shown as open blue circles, *ush2a^b1245^* mutants as filled red squares. The lines connect the mean values. Zeitgeber time 0 represents light onset at 4 dpf. Grey background represents the dark period between Zeitgeber time 14-24. No significant differences were detected between genotypes (unpaired t-test with Holm-Šídák multiple comparison correction).

### Circadian melatonin release from *ex vivo ush2a^b1245^* pineal glands remains intact

To assess whether usherin deficiency in the pineal gland affects circadian melatonin secretion, *ex vivo* pineal glands from *ush2a^b1245^* mutant zebrafish and wild-type sibling controls were superfused under constant darkness, and melatonin concentrations were measured across two circadian cycles. Both genotypes showed circadian melatonin release with nocturnal peaks and lower daytime levels. Peak timing did not differ between groups, and absolute melatonin concentrations did not differ significantly at any timepoint (False discovery rate-corrected unpaired t-tests). Group means, standard deviations, and p-values per timepoint are provided in supplementary table S3.

However, both groups showed substantial interindividual variability at each timepoint, with standard deviations often comparable to or exceeding the group means. To account for this interindividual variation, values were normalized per individual, and relative melatonin concentrations were used to fit a sine wave with nonzero baseline (Figure 5). The amplitude of the fitted melatonin rhythm was 0.490 in controls and 0.467 in *ush2a^b1245^* mutants (comparison of Fits, p = 0.826). The control pineal glands had a wavelength of 25.3 hours, compared to 24.3 hours in the *ush2a^b1245^* (p = 0.466). The phase shift was −0.0428 in controls and −0.483 in mutants (p = 0.204), and the baseline was 0.977 and 0.963, respectively (p = 0.841). The 95% confidence intervals for each parameter are provided in supplementary table S4.

**Figure 5.**
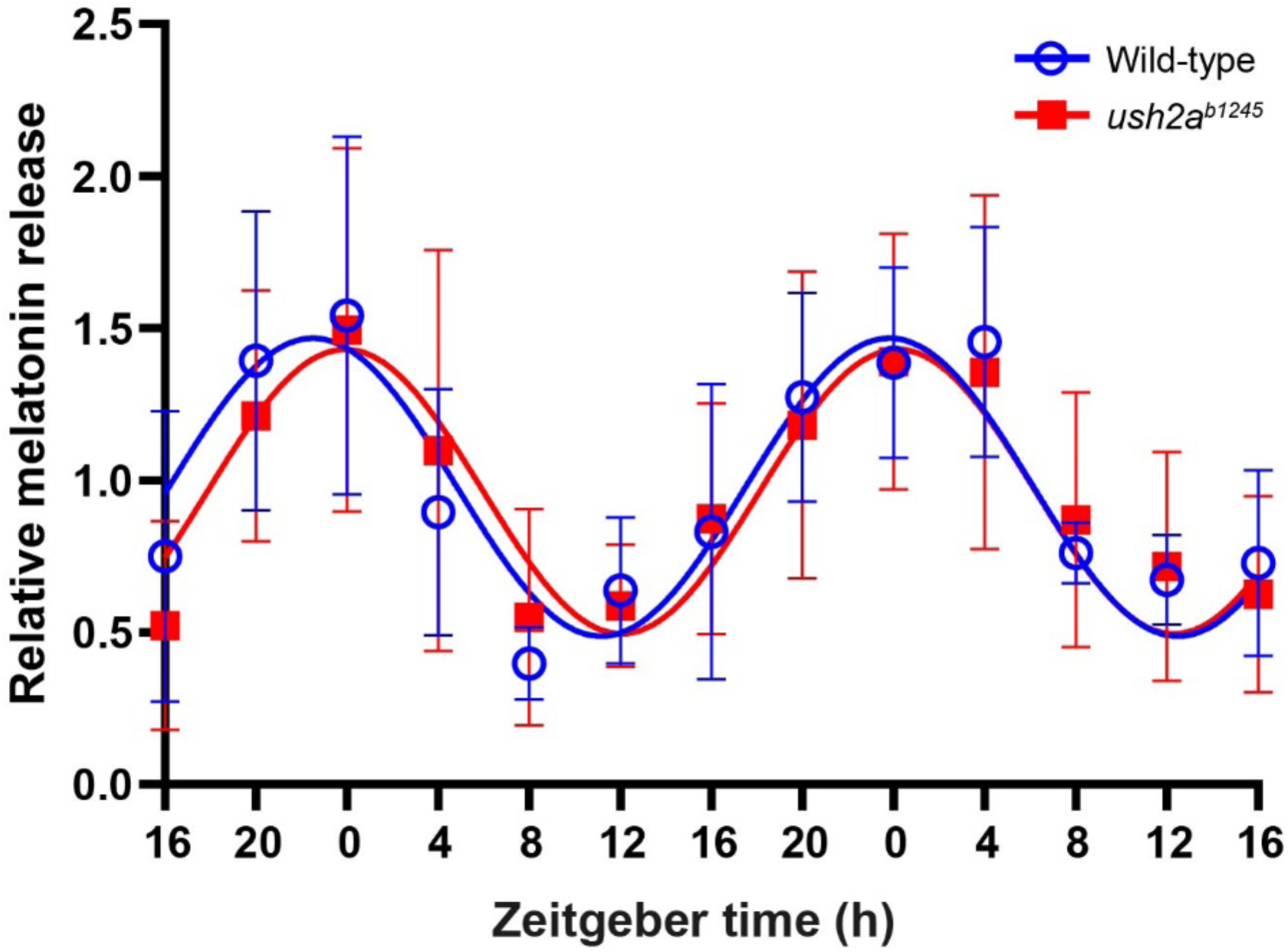
*Ex vivo* pineal glands of *ush2a^b1245^* mutants display sustained circadian melatonin release. Relative melatonin concentrations shown over time. Group mean of controls are shown in open blue circles, *ush2a^b1245^* mutants in filled red squares. Error bars represent the standard deviation. Coloured lines represent fitted circadian rhythms based on non-linear sine wave regression with a non-zero baseline. n = 6 per genotype per timepoint.

Together, these results indicate that the circadian rhythm of pineal melatonin release is not significantly altered in *ush2a^b1245^*mutants compared to controls within the timeframe of the experiment and with the used sample size. However, the substantially higher interindividual variability precludes firm conclusions about potential differences in absolute melatonin levels.

### *ush2a^b1245^* larvae retain circadian locomotor rhythms but show increased daytime sleep and prolonged sleep latency

To investigate whether loss of usherin affects the circadian sleep-wake behaviour, 24-hour video recordings of *ush2a^b1245^* and control larvae were performed using live-tracking of individual larvae under the conventional 14h:10h light/dark regime. The following behavioural states were quantified across the circadian cycle: wakefulness, three sleep substates with eye movement (QEM), each with different eye movement kinematics (QEM-1, QEM-2, QEM-3), and a sleep substate with no eye movement (QNEM) (Choudhary et al., 2023).

As shown in Figure 6A, the behavioural states of our control larvae followed the previously described circadian pattern by Choudhary et al. (2023): wakefulness predominated during the light phase, while QNEM, QEM-2, and QEM-3 were enriched during the dark phase. QEM-1 was the only sleep substate consistently displayed during the light period, and its probability was highest at 5 dpf, decreasing slightly at later stages.

**Figure 6.**
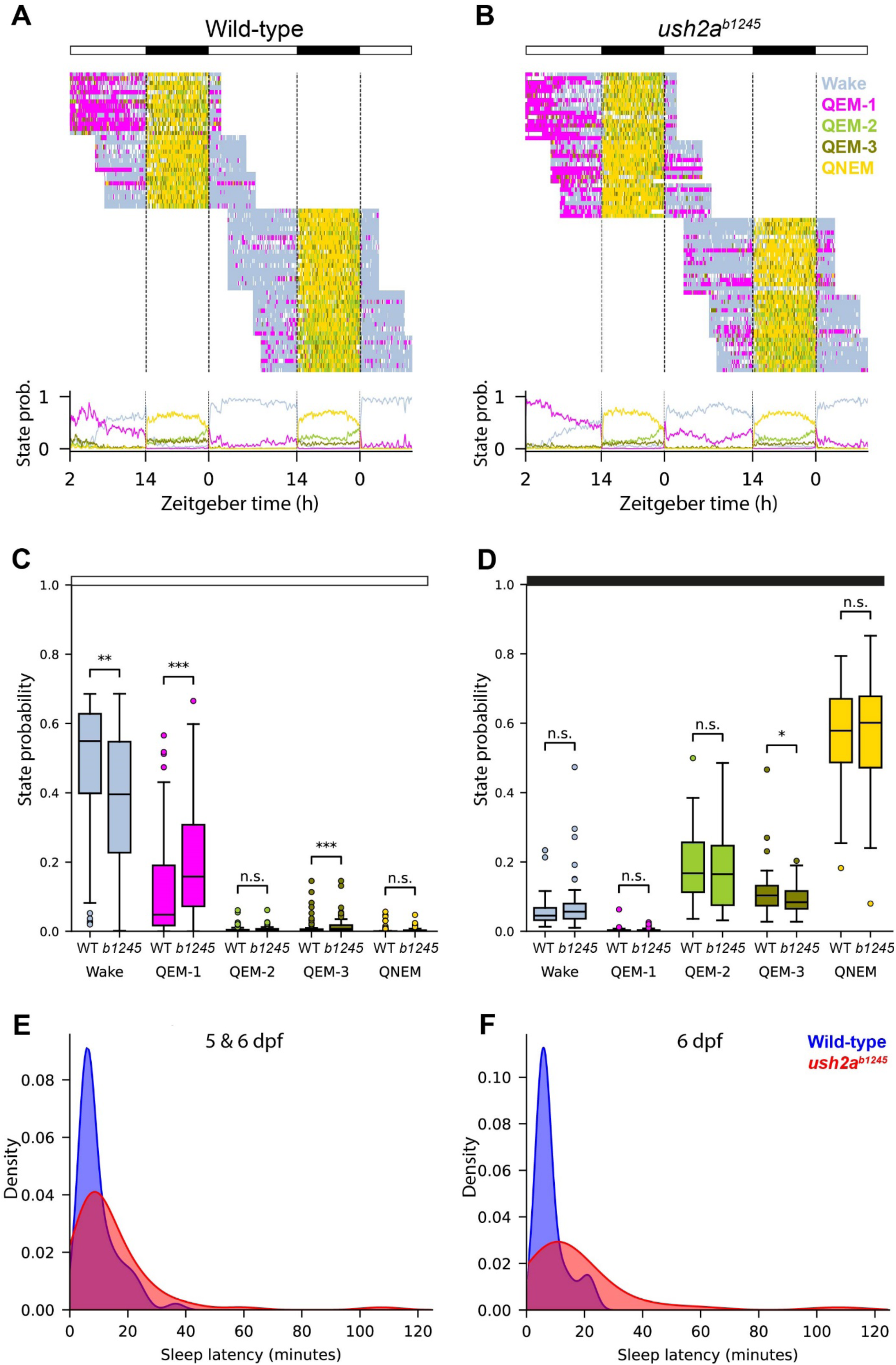
Circadian distribution of behavioural states and sleep latency in *ush2a^b1245^* mutants and control larvae. Behavioural states of control larvae, defined by locomotor activity and eye movement: Wake (light blue), QEM-1 (magenta), QEM-2 (green), QEM-3 (dark green), and QNEM (yellow). Horizontal bars above each graph indicate the light (ZT0-14) and dark (ZT14-24) periods. The data spans from 5-7dpf. **A)** Top: State classification raster plot of control larvae, where each row represents one individual (n = 66). Bottom: Corresponding temporal state probabilities. **B)** Same as in panel A, but for *ush2a^b1245^* mutants (n = 70). **C)** Comparison of daytime state probabilities between genotypes. Boxes show first to third quartile, in which the black horizontal line indicates the median. Whiskers extend to the most extreme data point within the 1.5 interquartile range. ∗ = p < 0.05; ∗∗ = p < 0.01; ∗∗∗ = p < 0.001 (Wilcoxon rank-sum test). **D)** Comparison of nighttime state probabilities between genotypes. **E)** Density plot showing the distribution of sleep latencies for control larvae (blue) and *ush2a^b1245^*mutants (red). Sleep latency was defined as the time from lights-off (ZT14) to the first transition into any sleep substate. Data from all ages tested (n = 66 control and n= 70 *ush2a^b1245^* larvae) **F)** Distribution of sleep latency at 6 dpf (n = 36 control and n= 36 *ush2a^b1245^* larvae).

In contrast, *ush2a^b1245^* mutants, displayed in Figure 6B, showed an altered temporal distribution of the behavioural states, with reduced daytime wakefulness and an increased QEM-1 probability during the light phase. Quantitative analysis of the behavioural states during daytime confirmed a significant reduction in daytime wake probability in *ush2a^b1245^* mutants (median = 0.396) compared to the controls (median = 0.549; p = 0.00525; Wilcoxon rank-sum test) (figure 6C). This reduction was accompanied by a marked increase in daytime QEM-1 probability (0.158 for *ush2a^b1245^* versus 0.0481 for controls; p = 0.000757). Additionally, a slight but highly significant increase in QEM-3 was observed (0.00741 in *ush2a^b1245^* versus 0.00204 in controls; p = 0.000375), although this state remained rare during the day in both genotypes. The daytime prevalence of QNEM and QEM-2 was low and did not differ significantly between groups.

As shown in Figure 5D, both genotypes exhibited the expected increase of QEM-2, QEM-3, and QNEM at night. However, *ush2a^b1245^* larvae showed a modest but significant decrease in QEM-3 probability compared to the control larvae (0.0841 for *ush2a^b1245^* versus 0.103 for controls; p = 0.0311), while wakefulness, QEM-1, QEM-2, and QNEM did not differ significantly. Together, these results show that *ush2a^b1245^* retain a circadian sleep-wake rhythm with sleep substates largely segregated between day and night, but they have a pronounced increase in daytime sleep driven primarily by elevated QEM-1 probability.

Next, the sleep latencies of both groups were compared, defined as the time taken to transition from wakefulness to any sleep substate after the onset of darkness at ZT14. As shown in figure 6E, *ush2a^b1245^* larvae exhibited significantly longer sleep latencies than controls across all ages tested, with a median of 8.68 minutes versus 6.61 minutes in the control larvae (p = 0.0216, Wilcoxon rank-sum test). Figure 5F shows that this difference was even more pronounced at 6 dpf, when the median latency in mutants was over 60% higher than in controls (10.08 versus 6.17 minutes for *ush2a^b1245^* and controls, respectively; p = 0.00279). In supplementary figure S5, zoom-ins on the state classification raster plots are given for the light-dark and dark-light transitions. When comparing the state probability of control larvae in figure S4A with the *ush2a^b1245^* larvae in S4B, the increase in daytime QEM-1 as well as the prolonged sleep latency is visible. Despite this delayed sleep latency, once sleep is established the mutant larvae seem to enter similar sleep states as controls. In Figure S4C and S4D it is shown that mutant larvae to transition from nighttime sleep (QEM-2, QEM-3 and QNEM) to normal daytime states (Wake and QEM-1) at the light onset (ZT0), although again with a higher QEM-1 probability than the controls.

In addition to the increase in both mean and median sleep latency, the sleep latency was also more variable between *ush2a^b1245^* larvae. Standard deviations were consistently higher in mutants, and Levene’s test confirmed significantly greater variance in sleep latency both when all ages were pooled (p = 0.0331) and at 6 dpf separately (p = 0.0132). Together, these results indicate that usherin loss delays and destabilizes the onset of nighttime sleep.

## DISCUSSION

The goal of this study was to investigate the biological basis underlying the sleep problems reported by USH2a patients, which we previously showed to occur independently of the severity of the vision loss. Using zebrafish models, we identified usherin expression in the pineal gland, a central regulator of sleep-wake cycles and circadian rhythmicity in all vertebrates. We furthermore observed significantly altered sleep behaviour in *ush2a* knock-out larvae. Compared to controls, usherin-deficient larvae displayed increased daytime QEM sleep and a prolonged sleep latency. Together, these findings indicate that usherin plays a role in sleep-wake behaviour regulation. No differences were detected in the expression of core circadian clock genes in larval zebrafish, and melatonin release from *ex vivo* cultured mutant and wild-type pineal glands displayed similar circadian phase and period. Our findings point to a previously unrecognized role for usherin in modulating sleep-wake behaviour, warranting further investigation into its underlying mechanisms.

Usherin is well established as a component of the retinal photoreceptor connecting cilium, where it is essential for providing structural integrity, vesicle docking, and protein trafficking. This study is the first to report usherin expression in the zebrafish pineal gland. Immunohistochemistry revealed that usherin, as well as USH2 protein complex members Adgrv1 and Whrn, are localised in close proximity of the basal body and cilium marker centrin in zebrafish pinealocytes. The presence of usherin in the pineal gland fits previously published transcriptomic datasets showing *USH2A* expression in mammalian pinealocytes (Bailey et al., 2009; Chang et al., 2020; Han et al., 2022). In vertebrate evolution, the pineal gland underwent marked morphological and functional changes. Whereas teleost pinealocytes are light-sensitive and possess a distinct photoreceptor morphology, mammalian pinealocytes lost direct photosensitivity and lack inner and outer segments (Ekström & Meissl, 2003; Falcón, 1999; Klein, 2006). The detection of usherin in porcine pineal gland tissue by Western blot demonstrates that the protein is still expressed in mammalian pinealocytes, even though these cells no longer possess the photoreceptive function and morphology as found in teleosts. The evolutionary conserved expression suggests that usherin serves a biologically relevant role in pinealocytes across species, independent of their photoreceptive morphology, and strengthens the translational relevance of our zebrafish findings to humans.

The behavioural data from *ush2a* knock-out larvae, showing elevated daytime sleep and significantly prolonged and more variable sleep latencies, provide strong evidence for an effect of usherin on sleep behaviour. Using the recently developed DASHER system (Choudhary et al., 2023), sleep-wake states were quantified with high spatiotemporal resolution. This study is the first translational application of this methodology and provides initial experimental support for its validity in modelling human sleep phenotypes. The observed aberrations in *ush2a* knock-out larvae are consistent with the findings from patient studies. The observed increased daytime sleep in *ush2a* zebrafish fits the fatigue and increased daytime sleepiness in patients (Hendricks et al., 2023). Additionally, the prolonged sleep latency and increased variability were also found in the patient actigraphy measurements (Hendricks et al., 2024). Importantly, the sleep phenotype of the *ush2a* mutant larvae occurred despite the relatively mild retinal phenotype (Dona et al., 2018), supporting a direct biological contribution of usherin loss to sleep dysregulation rather than an indirect effect of impaired light entrainment, again in accordance with the patient results (Hendricks et al., 2024; Hendricks et al., 2023).

To explore a potential role for usherin in circadian function, we examined clock gene expression in larvae and melatonin release in adult zebrafish. Expression of *aanat2*, *bmal1a*, *clock1a*, *cry1a*, *cry2a*, and *per2* was indistinguishable between mutants and controls under LD conditions. In contrast, *aanat2*, *clock1a* and *per2* knock-out larvae display marked alterations in circadian gene expression (Chen et al., 2023; Ren et al., 2017; Wang et al., 2015). These results indicate that the core transcriptional–translational feedback loop of the circadian clock remains intact in the *ush2a* mutants. The gene expression results align with the observations that *ush2a* larvae retain normal circadian timing of the different sleep stages. Subtle pineal-specific changes, however, cannot be excluded, as whole-larva qPCR could dilute tissue-specific aberrations due to the presence of light-entrainable clocks throughout the zebrafish body (Frøland Steindal & Whitmore, 2019). Additionally, expression profiles were not studied under prolonged constant darkness, as this does not reflect the clinical situation where patients maintain residual light perception.

*Ex vivo* superfusion of pineal glands under constant darkness showed that circadian melatonin release is preserved in *ush2a* mutants, with no detectable changes in phase and period. Although interindividual variability in release precludes firm conclusions about differences in absolute melatonin release, the presence of rhythmic release argues against a primary melatonin synthesis defect. This is consistent with the qPCR data showing unaltered expression of core circadian clock genes, together suggesting that the molecular circadian machinery remains intact in *ush2a* mutant larvae. Additionally, it aligns with the patient actigraphy data, where circadian sleep-wake rhythms remain intact (Hendricks et al., 2024). Nonetheless, it is important to consider the limitations of the superfusion method. The current sample size, large inter-individual variability, and relatively short measurement period preclude firm conclusions about absolute melatonin levels or subtle phase shifts between the groups. Moreover, the supplementation of the culture medium with tryptophan and 5-hydroxytryptophan, precursors of melatonin, may have masked potential upstream disruptions in the melatonin synthesis pathway (Emet et al., 2016). Therefore, while our findings argue against a major defect in core clock function or melatonin synthesis, additional studies are needed before such possibilities can be excluded. Measuring melatonin profiles in patients would be an important next step. Melatonin can be measured from saliva samples, providing a relatively non-invasive method. Importantly, such sampling can be performed at home without the need for a healthcare professional to be present, which reduces patient burden and allows assessment under normal conditions (Dermanowski et al., 2022). Such a patient study could clarify whether melatonin dynamics are altered in USH2a, and can inform about the direction of potential therapeutic interventions.

In this study, two independent *ush2a* mutant lines with different genetic backgrounds were examined. The *ush2a^b1245^* line (AB strain) expresses a truncated usherin protein with residual N-terminal immunoreactivity in pinealocytes, whereas usherin expression is not detected in the *ush2a^rmc1^*line (TL strain). Despite this difference, previous work demonstrated that the *ush2a^b1245^* line behaves as a functional null: the truncated protein is non-functional and leads to early-onset photoreceptor defects identical to a complete loss of usherin (Dona et al., 2018). Behavioural experiments were performed in *ush2a^b1245^* as the AB strain exhibits lower anxiety-induced behaviour and more pronounced light-dark responses than the TL strain (Egan et al., 2009; van den Bos et al., 2017), thereby providing a more robust baseline against which subtle sleep phenotypes can be detected. Since the truncated usherin protein is non-functional, and *ush2a^rmc1^*show comparable circadian clock gene expression patterns as observed in *ush2a^b1245^*, it is expected that the sleep phenotype identified in *ush2a^b1245^* can be generalised across *ush2a* knock-out zebrafish models that have been generated against different backgrounds.

Further work is needed to identify the molecular mechanisms underlying the sleep aberrations. Transcriptomic profiling of dissected pineal glands from *ush2a* mutants and controls at multiple Zeitgeber times could provide relevant insights into the molecular pathways linking usherin dysfunction to altered sleep-wake behaviour, yet this is technically challenging due to the need to dissect zebrafish in near-complete darkness. Such an approach would provide pineal gland-specific data, allowing detection of aberrations that remain undetectable in whole-larva samples. And most importantly, it allows unbiased identification of affected pathways, potentially revealing molecular changes not captured in this study. The eventual development of diurnal mammalian USH2(a) models such as pig and non-human primate would enable cross-species validation and contribute to the translatability of the findings. This is particularly relevant because mammalian pinealocytes have diverged substantially from their photoreceptive ancestors. Understanding why usherin is conserved in morphologically divergent pineal glands, and how its loss can impact sleep behaviour across distinct species, remains an intriguing knowledge gap with both fundamental and clinical relevance. Beyond melatonin synthesis, one plausible downstream mechanism could involve other altered neuroendocrine signalling. Usher proteins are present in the synaptic ribbons of hair cells and photoreceptors (Kremer et al., 2006), and mammalian pinealocytes likewise contain synaptic ribbons that mediate secretory functions (Al-Hussain, 2006; Spiwoks-Becker et al., 2008). Loss of usherin might alter neuronal or endocrine output from the pineal gland, affecting the release of signalling molecules that modulate arousal and sleep (Smith & Mong, 2019).

In conclusion, this study shows that usherin localises to the pineal gland and that the loss of this protein leads to measurable aberrations in sleep behaviour. These findings establish the first molecular link between usherin and circadian sleep behaviour in a vertebrate model. The conservation of pineal usherin expression in higher vertebrates further supports the translational relevance of these findings and provides a biological basis for the sleep disturbances reported in USH2a patients. As the underlying mechanisms remain partially unresolved, follow-up studies such as pineal gland RNA sequencing and patient melatonin profiling will be essential to clarify how usherin contributes to sleep regulation. Altogether, this work lays the groundwork for future development of therapeutic strategies aimed at improving sleep - and ultimately quality of life - of USH2a patients.

## Acknowledgements

A special thanks Prof. Dr. Monte Westerfield and Dr. Jennifer Phillips from the University of Oregon for generating and providing the *ush2a^b1245^* zebrafish line, and to the team of Prof. Dr. Nikolai Klymiuk from the Technical University of Munich and Prof. Dr. Jan Motlik from the PIGMOD Center in Liběchov for the dissection of the pig pineal gland. This study was supported by Stichting Ushersyndroom, Dr C.J. Vaillantfonds, Landelijke Stichting voor Blinden en Slechtzienden, Steunfonds Uitzicht (Beheer’t Schild), Algemene Nederlandse Vereniging ter Voorkoming van Blindheid (UZ2020-16), de Gelderse Blindenstichting, and Radboud Institute for Biological and Environmental Sciences (Radboud University).

## SUPPLEMENTARY TABLES AND FIGURES

**Supplementary Table S1.**
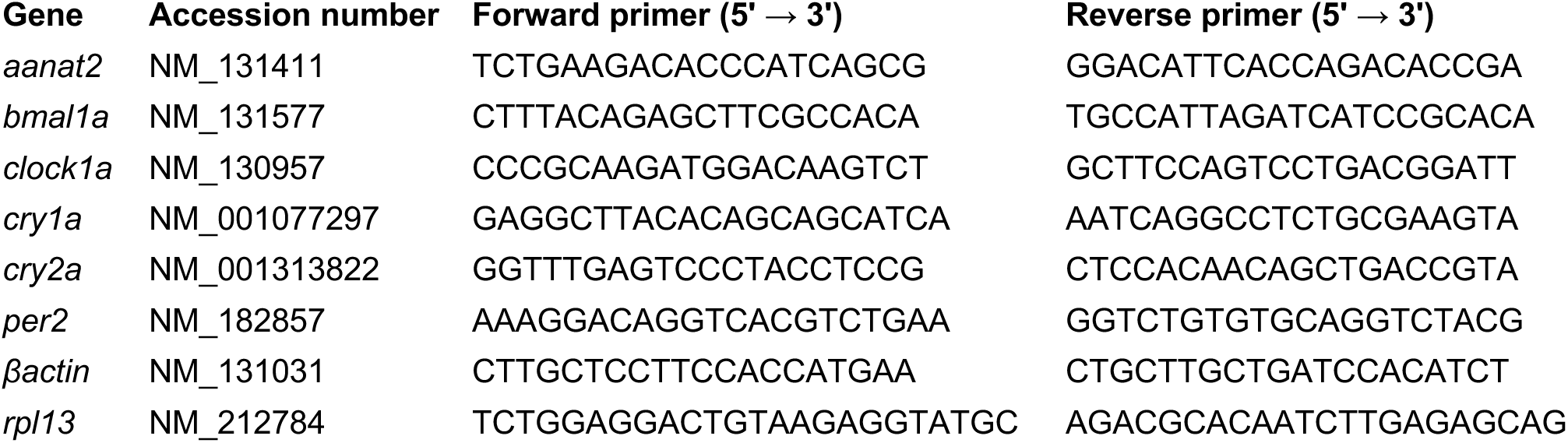
Primer sequences used for qPCR analysis of circadian clock genes. βactin and rpl13 served as housekeeping genes. Accession numbers refer to Ensembl transcript identifiers.

**Supplementary Table S2.** Amino acids in culture medium. Final concentration in mM shown. All amino acids were first dissolved in 100x stock solutions and then added to the culture medium.

| **Amino acid** | **Concentration (mM)** | **Product number** |
| --- | --- | --- |
| L-arginine | 0.14 | A6969, Sigma-Aldrich |
| L-cystine | 0.09 | C8755, Sigma-Aldrich |
| L-glutamine | 0.84 | 35050061, Gibco |
| Glycine | 0.25 | G7126, Sigma-Aldrich |
| L-histidine | 0.16 | H5659, Sigma-Aldrich |
| L-isoleucine | 0.30 | I7403, Sigma-Aldrich |
| L-leucine | 0.30 | L8000, Sigma-Aldrich |
| L-lysine | 0.33 | L8662, Sigma-Aldrich |
| L-methionine | 0.08 | M5308, Sigma-Aldrich |
| L-phenylalanine | 0.16 | P5482, Sigma-Aldrich |
| L-threonine | 0.33 | T8625, Sigma-Aldrich |
| L-tryptophan | 0.04 | T8941, Sigma-Aldrich |
| L-tyrosine | 0.16 | T8566, Sigma-Aldrich |
| L-valine | 0.33 | V0500, Sigma-Aldrich |

**Supplementary Table S3.** Melatonin levels across time points in control and *ush2a^b1245^* pineal glands. Mean ± SD melatonin concentration (pg/h) per time point for each genotype (n = 6 per group). Statistical significance assessed by unpaired t-tests with false discovery rate correction for multiple comparisons. ZT = Zeitgeber time.

| **ZT** | **Control**<br>**Mean ± SD** | ***ush2a*<sup>b1245</sup>**<br>**Mean ± SD** | **p-value** |
| --- | --- | --- | --- |
| 14-16 | 258 ± 307 | 135 ± 188 | 0.424 |
| 18-20 | 443 ± 429 | 289 ± 322 | 0.498 |
| 22-0 | 446 ± 443 | 297 ± 334 | 0.527 |
| 2-4 | 241 ± 244 | 152 ± 185 | 0.496 |
| 6-8 | 112 ± 102 | 81.8 ± 105 | 0.623 |
| 10-12 | 227 ± 201 | 131 ± 167 | 0.389 |
| 14-16 | 350 ± 355 | 266 ± 322 | 0.678 |
| 18-20 | 473.4 ± 423 | 382 ± 436 | 0.719 |
| 22-0 | 465.6 ± 396 | 448 ± 565 | 0.952 |
| 2-4 | 474.2 ± 418 | 436 ± 685 | 0.910 |

**Supplementary Table S4.** Confidence intervals (CI) for best-fit parameters of melatonin rhythm. 95% CI for amplitude, wavelength, phaseshift, and baseline of the fitted circadian melatonin rhythm in control and *ush2a^b1245^*pineal glands. n = 6 per genotype per timepoint.

| **Best-fit value** | **Control 95% CI** | ***ush2a*<sup>b1245</sup> 95% CI** |
| --- | --- | --- |
| Amplitude | 0.359 - 0.621 | 0.308 - 0.627 |
| Wavelength | 23.6 - 27.6 | 22.2 - 26.9 |
| Phaseshift | -0.481 - 0.435 | -1.05 - 0.0735 |
| Baseline | 0.885 - 1.07 | 0.840 - 1.09 |

**Supplementary Figure S1.**
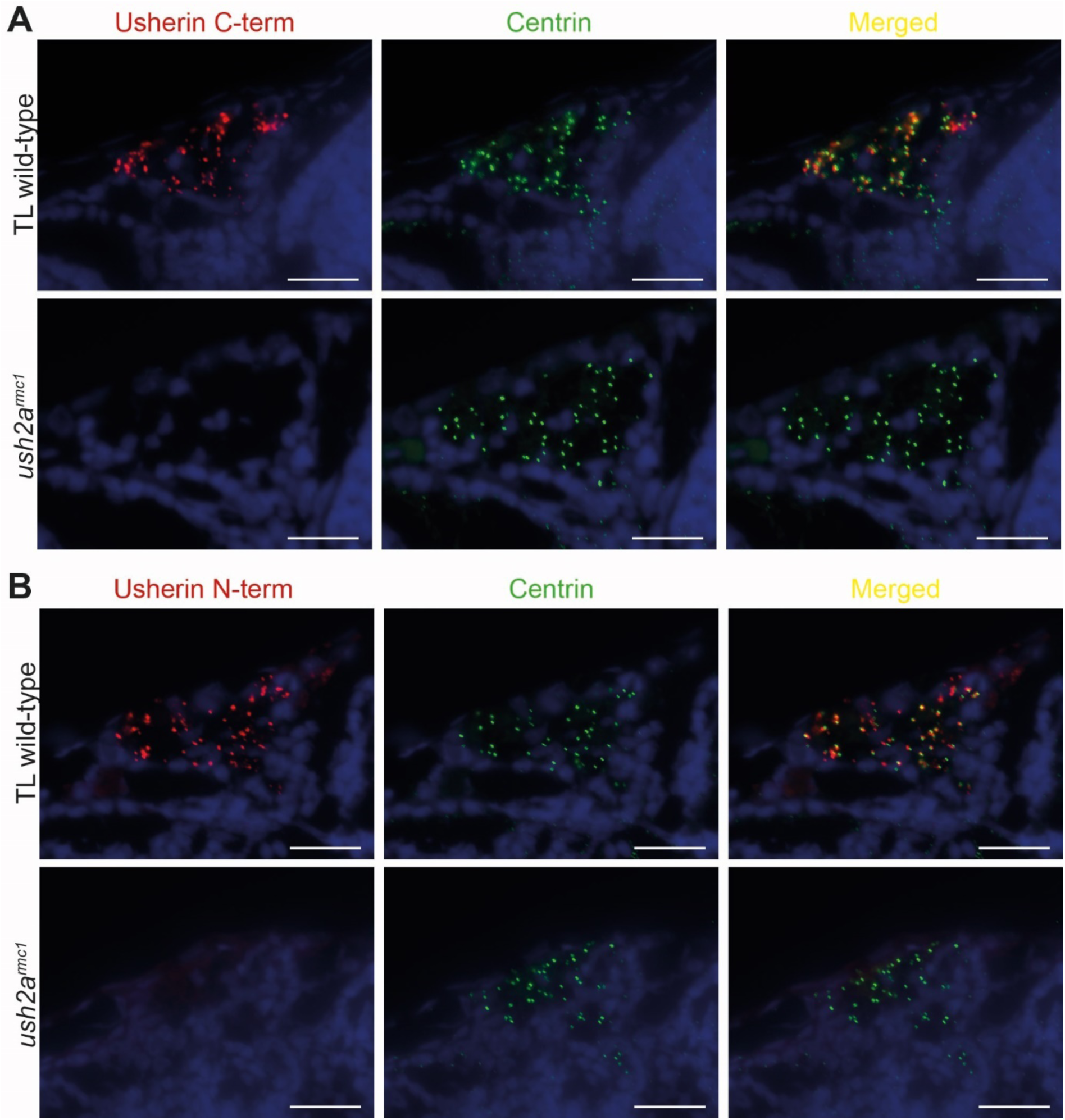
Absence of usherin in the pineal gland of *ush2a^rmc1^* zebrafish larvae. **A)** Immunofluorescence labelling of the pineal gland in TL wild-type and *ush2a^rmc1^* mutant larvae showing DAPI (blue), usherin C-terminus (red), and centrin (green). **B)** Corresponding immunofluorescence images with usherin N-terminus. In the TL wild-type pineal gland, usherin co-localises with the connecting cilium marker centrin. No usherin staining was detected with either antibody in *ush2a^rmc1^*mutants, which carry an early truncating mutation in the *ush2a* gene. Scale bar: 20 µm.

**Supplementary Figure S2.**
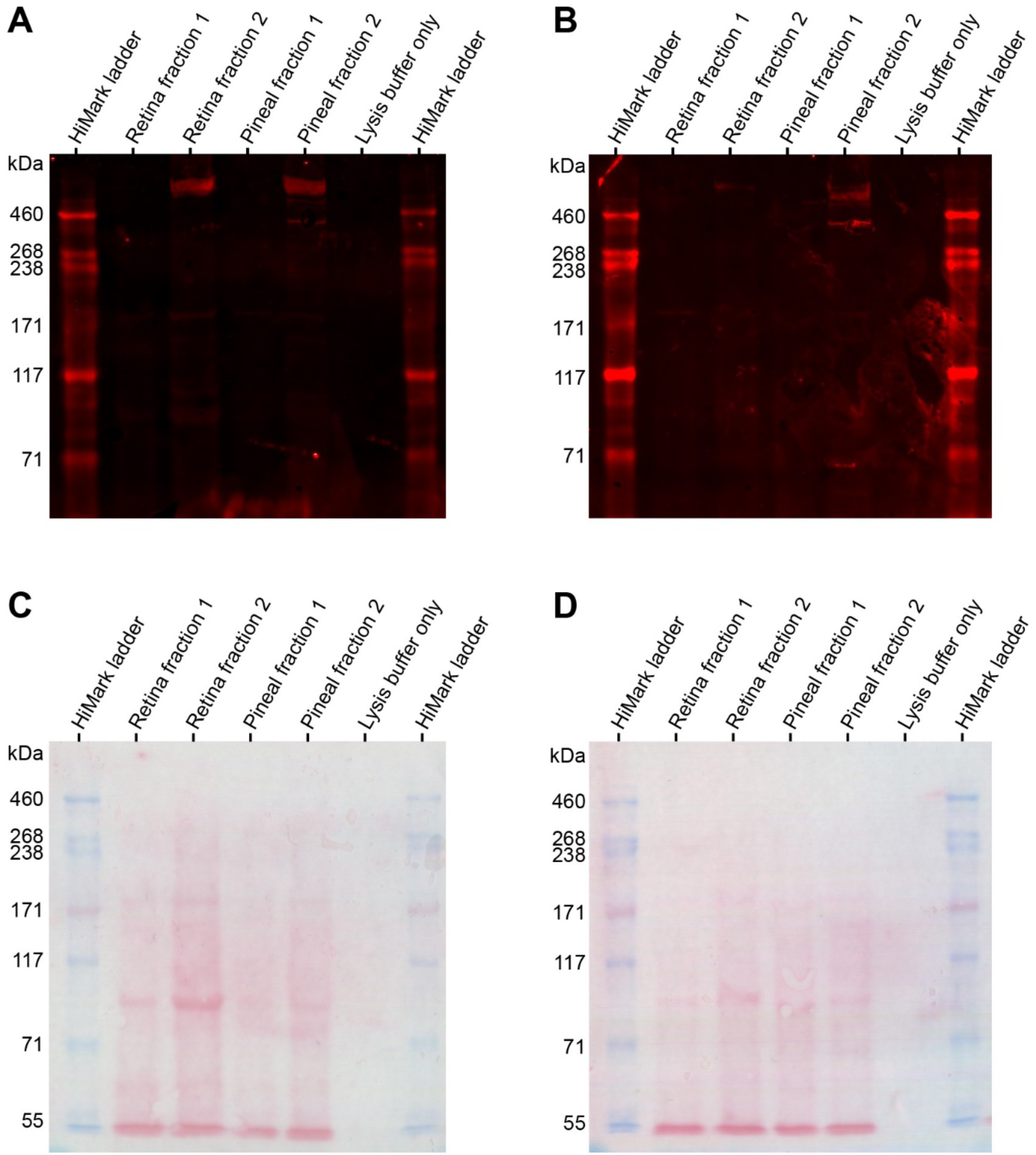
Full western blot images and biological replicate for detection of usherin protein in pig retina and pineal gland. **A-B)** Western blot against usherin, showing a high-molecular-weight band of ∼570 kDa in the second fraction of both retinal and pineal gland lysates in two independent experiments. HiMark ladder indicates molecular weight standards. **C-D)** Corresponding Ponceau S staining of the membranes, confirming protein transfer and comparable loading across lanes.

**Supplementary Figure S3.**
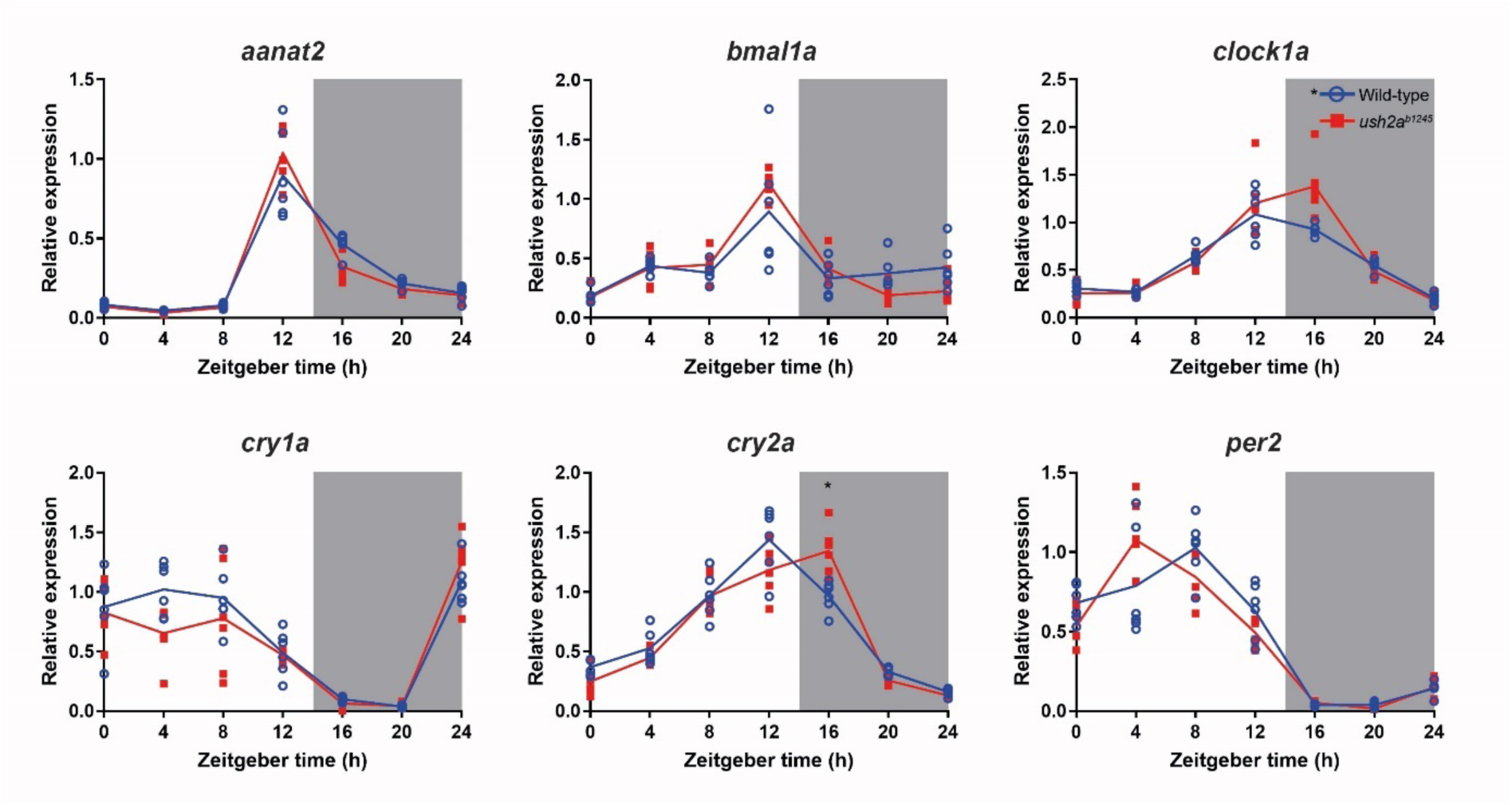
Relative gene expression of clock genes in 4 dpf larvae. Expression measured over a 24h period at 4-hour intervals. Individual measurements are shown as dots (n=6). TL wild-type larvae are shown as open blue circles, *ush2a^rmc1^* mutants as filled red squares. The lines connect the mean values. Zeitgeber time 0 represents light onset at 4 dpf. Grey background represents the dark period between ZT14-24. *Ush2a^rmc1^*mutants show the same trends as controls, with minor differences for *clock1a* and *cry2a* at ZT16. Unpaired t-test with Holm-Šídák multiple comparison correction, * = p <0.05.

**Supplementary Figure S5.**
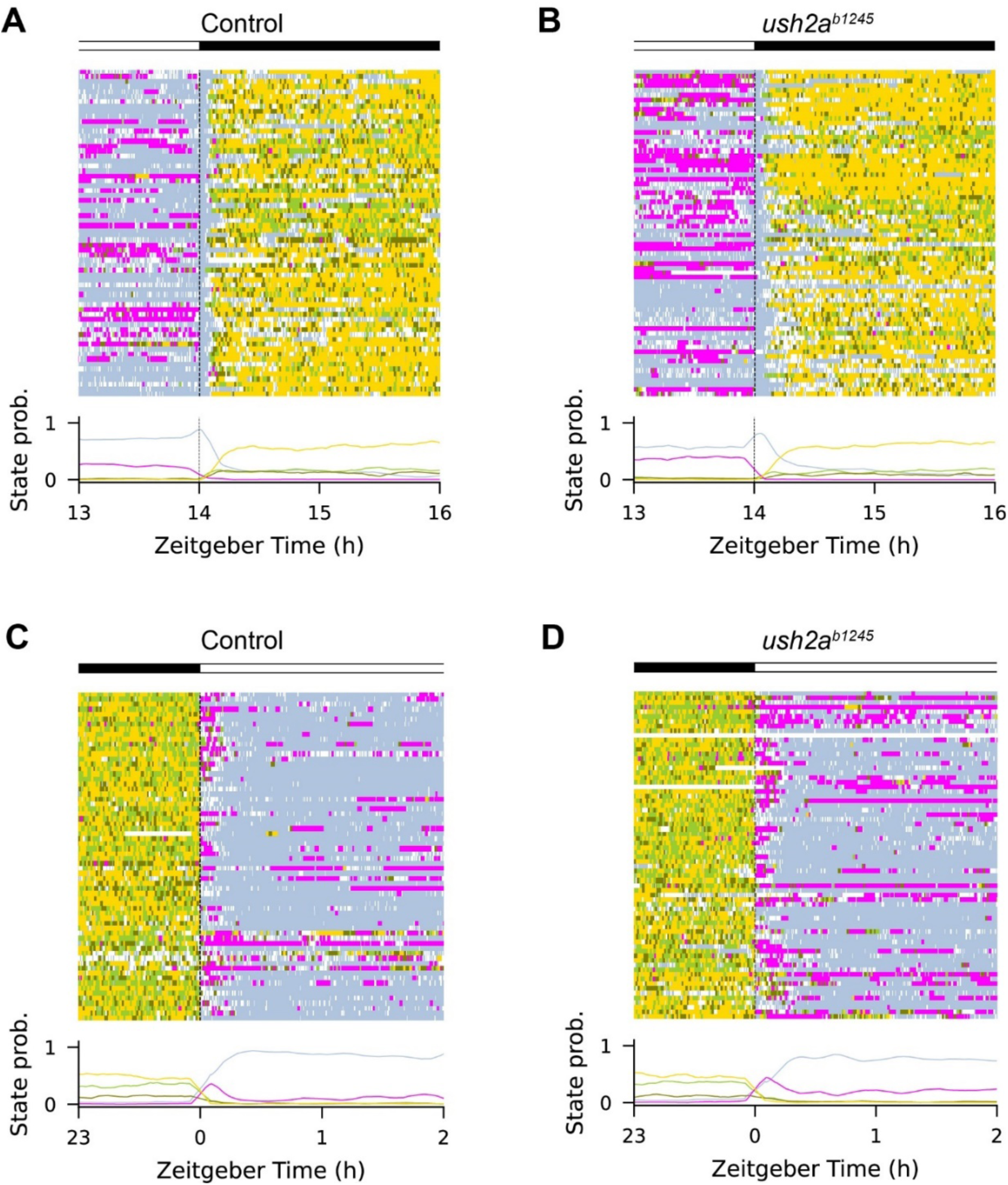
Zoom-in on behavioural states during light-dark and dark-light transitions. Behavioural states of control larvae, defined by locomotor activity and eye movement: Wake (light blue), QEM-1 (magenta), QEM-2 (green), QEM-3 (dark green), and QNEM (yellow). Horizontal bars above each graph indicate the light (white) and dark periods (black). Top: raster plots showing individual larvae. Bottom: corresponding temporal state-probability profiles. **A)** Control larvae around lights-off (ZT 13-16, n = 66). **B)** *ush2a^b1245^* mutants around lights-off (n = 70). **C)** Control larvae around lights-on (ZT23-2). **D)** *ush2a^b1245^* mutants around lights-on.

## References

Al-Hussain, S. (2006). The pinealocytes of the human pineal gland: a light and electron microscopic study. Folia morphologica, 65(3), 181–187.

Aleström, P., D’Angelo, L., Midtlyng, P. J., Schorderet, D. F., Schulte-Merker, S., Sohm, F., & Warner, S. (2019). Zebrafish: Housing and husbandry recommendations. Laboratory Animals, 54(3), 213–224. 10.1177/0023677219869037

Appelbaum, L., Wang, G. X., Maro, G. S., Mori, R., Tovin, A., Marin, W., Yokogawa, T., Kawakami, K., Smith, S. J., & Gothilf, Y. J. P. o. t. N. A. o. S. (2009). Sleep–wake regulation and hypocretin–melatonin interaction in zebrafish. 106(51), 21942–21947.

Bahena, P., Daftarian, N., Maroofian, R., Linares, P., Villalobos, D., Mirrahimi, M., Rad, A., Doll, J., Hofrichter, M. A., & Koparir, A. (2022). Unraveling the genetic complexities of combined retinal dystrophy and hearing impairment. Human Genetics, 141(3), 785–803.

Bailey, M. J., Coon, S. L., Carter, D. A., Humphries, A., Kim, J.-s., Shi, Q., Gaildrat, P., Morin, F., Ganguly, S., & Hogenesch, J. B. (2009). Night/Day Changes in Pineal Expression of> 600 Genes CENTRAL ROLE OF ADRENERGIC/cAMP SIGNALING. Journal of Biological Chemistry, 284(12), 7606–7622. https://www.ncbi.nlm.nih.gov/pmc/articles/PMC2658055/pdf/7606.pdf

Berson, D. M., Dunn, F. A., & Takao, M. (2002). Phototransduction by retinal ganglion cells that set the circadian clock [Article]. Science, 295(5557), 1070–1073. 10.1126/science.1067262

Besharse, J. C., & Dunis, D. A. (1983). Rod photoreceptor disc shedding in eye cups: relationship to bicarbonate and amino acids. Experimental eye research, 36(4), 567–579.

Cahill, G., & Besharse, J. (1991). Resetting the circadian clock in cultured Xenopus eyecups: regulation of retinal melatonin rhythms by light and D2 dopamine receptors. The Journal of neuroscience: the official journal of the Society for Neuroscience, 11(10), 2959. http://www.jneurosci.org/content/jneuro/11/10/2959.full.pdf

Cahill, G. M. (1996). Circadian regulation of melatonin production in cultured zebrafish pineal and retina. Brain research, 708(1-2), 177–181.

Casadei, R., Pelleri, M. C., Vitale, L., Facchin, F., Lenzi, L., Canaider, S., Strippoli, P., & Frabetti, F. (2011). Identification of housekeeping genes suitable for gene expression analysis in the zebrafish. Gene Expression Patterns, 11(3), 271–276.

Chang, E., Fu, C., Coon, S. L., Alon, S., Bozinoski, M., Breymaier, M., Bustos, D. M., Clokie, S. J., Gothilf, Y., Esnault, C., Michael Iuvone, P., Mason, C. E., Ochocinska, M. J., Tovin, A., Wang, C., Xu, P., Zhu, J., Dale, R., & Klein, D. C. (2020). Resource: A multi-species multi-timepoint transcriptome database and webpage for the pineal gland and retina. Journal of pineal research, 69(3), e12673. 10.1111/jpi.12673

Chen, A.-q., Xue, M., Qiu, C.-z., Zhang, H.-y., Zhou, R., Zhang, L., Yin, Z.-j., & Ren, D.-l. (2023). Circadian clock1a coordinates neutrophil recruitment via nfe212a/duox-reactive oxygen species pathway in zebrafish. Cell Reports, 42(10), 113179. 10.1016/j.celrep.2023.113179

Choudhary, V., Heller, C. R., Aimon, S., de Sardenberg Schmid, L., Robson, D. N., & Li, J. M. (2023). Neural and behavioral organization of rapid eye movement sleep in zebrafish. bioRxiv, 2023.2008. 2028.555077.

Copper, J. E., Budgeon, L. R., Foutz, C. A., van Rossum, D. B., Vanselow, D. J., Hubley, M. J., Clark, D. P., Mandrell, D. T., & Cheng, K. C. (2018). Comparative analysis of fixation and embedding techniques for optimized histological preparation of zebrafish. Comparative Biochemistry and Physiology Part C: Toxicology & Pharmacology, 208, 38–46.

Dermanowski, M. M., Hejduk, A., Kuczyńska, J., Wichniak, A., Urbańska, A., & Mierzejewski, P. (2022). Assessment of dim light melatonin onset based on plasma and saliva samples. Chronobiology International, 39(5), 626–635.

Dona, M., Slijkerman, R., Lerner, K., Broekman, S., Wegner, J., Howat, T., Peters, T., Hetterschijt, L., Boon, N., de Vrieze, E., & van Wijk, E. (2018). Usherin defects lead to early-onset retinal dysfunction in zebrafish. 173, 148–159. https://www.sciencedirect.com/science/article/pii/S0014483518302458?via%3Dihub

Egan, R. J., Bergner, C. L., Hart, P. C., Cachat, J. M., Canavello, P. R., Elegante, M. F., Elkhayat, S. I., Bartels, B. K., Tien, A. K., Tien, D. H., Mohnot, S., Beeson, E., Glasgow, E., Amri, H., Zukowska, Z., & Kalueff, A. V. (2009). Understanding behavioral and physiological phenotypes of stress and anxiety in zebrafish. Behav Brain Res, 205(1), 38–44. 10.1016/j.bbr.2009.06.022

Ehn, M., Wahlqvist, M., Möller, C., & Anderzén-Carlsson, A. (2020). The lived experiences of work and health of people living with deaf-blindness due to Usher syndrome type 2. International Journal of Qualitative Studies on Health and Well-being, 15(1), 1846671. https://www.ncbi.nlm.nih.gov/pmc/articles/PMC7734013/pdf/ZQHW_15_1846671.pdf

Ekström, P., & Meissl, H. (2003). Evolution of photosensory pineal organs in new light: the fate of neuroendocrine photoreceptors. Philosophical Transactions of the Royal Society of London. Series B: Biological Sciences, 358(1438), 1679–1700. https://www.ncbi.nlm.nih.gov/pmc/articles/PMC1693265/pdf/14561326.pdf

Emet, M., Ozcan, H., Ozel, L., Yayla, M., Halici, Z., & Hacimuftuoglu, A. J. T. E. j. o. m. (2016). A review of melatonin, its receptors and drugs. 48(2), 135.

Esquiva, G., Lax, P., & Cuenca, N. (2013). Impairment of intrinsically photosensitive retinal ganglion cells associated with late stages of retinal degeneration. Investigative ophthalmology & visual science, 54(7), 4605–4618.

Falcón, J. (1999). Cellular circadian clocks in the pineal. Progress in neurobiology, 58(2), 121–162. https://www.sciencedirect.com/science/article/pii/S0301008298000781?via%3Dihub

Frenzel, H., Bohlender, J., Pinsker, K., Wohlleben, B., Tank, J., Lechner, S. G., Schiska, D., Jaijo, T., Rüschendorf, F., & Saar, K. (2012). A genetic basis for mechanosensory traits in humans. PLoS biology, 10(5), e1001318. https://www.ncbi.nlm.nih.gov/pmc/articles/PMC3341339/pdf/pbio.1001318.pdf

Frøland Steindal, I. A., & Whitmore, D. (2019). Circadian Clocks in Fish-What Have We Learned so far? Biology (Basel), 8(1). 10.3390/biology8010017

Han, L., Wei, X., Liu, C., Volpe, G., Zhuang, Z., Zou, X., Wang, Z., Pan, T., Yuan, Y., & Zhang, X. (2022). Cell transcriptomic atlas of the non-human primate Macaca fascicularis. Nature, 604(7907), 723–731.

Hartley, S., Dauvilliers, Y., & Quera-Salva, M.-A. (2018). Circadian rhythm disturbances in the blind. Current neurology and neuroscience reports, 18(10), 65. https://link.springer.com/article/10.1007/s11910-018-0876-9

Hendricks, J. M., Metz, J. R., Boss, H. M., Collin, R. W., de Vrieze, E., & van Wijk, E. (2024). Actigraphy-based assessment of circadian rhythmicity and sleep in patients with Usher syndrome type 2a: A case–control study. Journal of sleep research, e14456.

Hendricks, J. M., Metz, J. R., Velde, H. M., Weeda, J., Hartgers, F., Yzer, S., Hoyng, C. B., Pennings, R. J., Collin, R. W., & Boss, M. H. (2023). Evaluation of Sleep Quality and Fatigue in Patients with Usher Syndrome Type 2a. Ophthalmology Science, 3(4), 100323.

Jansen, F., Kalbe, B., Scholz, P., Mikosz, M., Wunderlich, K. A., Kurtenbach, S., Nagel-Wolfrum, K., Wolfrum, U., Hatt, H., & Osterloh, S. (2016). Impact of the Usher syndrome on olfaction. Human Molecular Genetics, 25(3), 524–533.

Karuntu, J. S., Almushattat, H., Nguyen, X. T., Plomp, A. S., Wanders, R. J. A., Hoyng, C. B., van Schooneveld, M. J., Schalij-Delfos, N. E., Brands, M. M., Leroy, B. P., van Karnebeek, C. D. M., Bergen, A. A., van Genderen, M. M., & Boon, C. J. F. (2025). Syndromic retinitis pigmentosa. Prog Retin Eye Res, 107, 101324. 10.1016/j.preteyeres.2024.101324

Katoh, K., Rozewicki, J., & Yamada, K. D. (2017). MAFFT online service: multiple sequence alignment, interactive sequence choice and visualization. Briefings in Bioinformatics, 20(4), 1160–1166. 10.1093/bib/bbx108

Klein, D. C. (2006). Evolution of the vertebrate pineal gland: the AANAT hypothesis. 23(1-2), 5–20. https://www.tandfonline.com/doi/pdf/10.1080/07420520500545839?needAccess=true

Kremer, H., van Wijk, E., Märker, T., Wolfrum, U., & Roepman, R. (2006). Usher syndrome: molecular links of pathogenesis, proteins and pathways. Human Molecular Genetics, 15(suppl_2), R262–R270.

Laurà, R., Magnoli, D., Zichichi, R., Guerrera, M. C., De Carlos, F., Suárez, A. Á., Abbate, F., Ciriaco, E., Vega, J. A., & Germanà, A. (2012). The photoreceptive cells of the pineal gland in adult zebrafish (Danio rerio). Microscopy research and technique, 75(3), 359–366. https://analyticalsciencejournals.onlinelibrary.wiley.com/doi/full/10.1002/jemt.21064

LeGates, T. A., Fernandez, D. C., & Hattar, S. (2014). Light as a central modulator of circadian rhythms, sleep and affect. Nature Reviews Neuroscience, 15(7), 443–454.

Letunic, I., & Bork, P. (2024). Interactive Tree of Life (iTOL) v6: recent updates to the phylogenetic tree display and annotation tool. Nucleic acids research, 52(W1), W78–W82. 10.1093/nar/gkae268

Liu, X., Bulgakov, O. V., Darrow, K. N., Pawlyk, B., Adamian, M., Liberman, M. C., & Li, T. (2007). Usherin is required for maintenance of retinal photoreceptors and normal development of cochlear hair cells. Proceedings of the National Academy of Sciences, 104(11), 4413–4418. doi:10.1073/pnas.0610950104

Livne, Z. B.-M., Alon, S., Vallone, D., Bayleyen, Y., Tovin, A., Shainer, I., Nisembaum, L. G., Aviram, I., Smadja-Storz, S., & Fuentes, M. J. P. g. (2016). Genetically blocking the zebrafish pineal clock affects circadian behavior. 12(11), e1006445. https://journals.plos.org/plosgenetics/article?id=10.1371/journal.pgen.1006445

Machado, T., Cortinhal, T., Carvalho, A. L., Teixeira-Marques, F., Silva, R., Murta, J., & Marques, J. P. (2025). Unraveling the genetic spectrum of inherited deaf-blindness in Portugal. Orphanet Journal of Rare Diseases, 20(1), 22.

Mathur, P., & Yang, J. (2015). Usher syndrome: hearing loss, retinal degeneration and associated abnormalities. Biochimica et Biophysica Acta (BBA)-Molecular Basis of Disease, 1852(3), 406–420. https://www.ncbi.nlm.nih.gov/pmc/articles/PMC4312720/pdf/nihms646774.pdf

McCurley, A. T., & Callard, G. V. (2008). Characterization of housekeeping genes in zebrafish: male-female differences and effects of tissue type, developmental stage and chemical treatment. BMC molecular biology, 9(1), 102.

Millán, J. M., Aller, E., Jaijo, T., Blanco-Kelly, F., Gimenez-Pardo, A., & Ayuso, C. (2010). An update on the genetics of usher syndrome. Journal of ophthalmology, 2011.

Pierrache, L. H., Hartel, B. P., Van Wijk, E., Meester-Smoor, M. A., Cremers, F. P., De Baere, E., De Zaeytijd, J., Van Schooneveld, M. J., Cremers, C. W., & Dagnelie, G. (2016). Visual prognosis in USH2A-associated retinitis pigmentosa is worse for patients with Usher syndrome type IIa than for those with nonsyndromic retinitis pigmentosa. Ophthalmology, 123(5), 1151–1160.

Ramos, J. N., Ribeiro, J. C., Pereira, A. C., Ferreira, S., Duarte, I. C., & Castelo-Branco, M. J. N. C. (2019). Evidence for impaired olfactory function and structural brain integrity in a disorder of cilia function, USHER syndrome. Neuroimage Clin, 22, 101757. 10.1016/j.nicl.2019.101757

Ren, D.-l., Ji, C., Wang, X.-B., Wang, H., & Hu, B. J. S. r. (2017). Endogenous melatonin promotes rhythmic recruitment of neutrophils toward an injury in zebrafish. 7(1), 4696.

Reurink, J., de Vrieze, E., Li, C. H., van Berkel, E., Broekman, S., Aben, M., Peters, T., Oostrik, J., Neveling, K., Venselaar, H., & van Wijk, E. (2022). Scrutinizing pathogenicity of the USH2A c. 2276 G> T; p.(Cys759Phe) variant. npj Genomic Medicine, 7(1), 1–9.

Riazuddin, S., Belyantseva, I. A., Giese, A. P., Lee, K., Indzhykulian, A. A., Nandamuri, S. P., Yousaf, R., Sinha, G. P., Lee, S., & Terrell, D. (2025). Author Correction: Alterations of the CIB2 calcium-and integrin-binding protein cause Usher syndrome type 1J and nonsyndromic deafness DFNB48. Nature genetics, 1–3.

Rihel, J., Prober, D. A., & Schier, A. F. (2010). Monitoring sleep and arousal in zebrafish. In Methods in cell biology (Vol. 100, pp. 281–294). Elsevier.

Sanjurjo-Soriano, C., Jimenez-Medina, C., Erkilic, N., Cappellino, L., Lefevre, A., Nagel-Wolfrum, K., Wolfrum, U., Van Wijk, E., Roux, A.-F., & Meunier, I. (2023). USH2A variants causing retinitis pigmentosa or Usher syndrome provoke differential retinal phenotypes in disease-specific organoids. Human Genetics and Genomics Advances, 4(4).

Schwaller, F., Bégay, V., García-García, G., Taberner, F. J., Moshourab, R., McDonald, B., Docter, T., Kühnemund, J., Ojeda-Alonso, J., Paricio-Montesinos, R., Lechner, S. G., Poulet, J. F. A., Millan, J. M., & Lewin, G. R. (2021). USH2A is a Meissner’s corpuscle protein necessary for normal vibration sensing in mice and humans. Nat Neurosci, 24(1), 74–81. 10.1038/s41593-020-00751-y

Slijkerman, R. W., Kremer, H., & van Wijk, E. (2017). Molecular Genetics of Usher Syndrome: Current State of Understanding. 1–12.

Slijkerman, R. W., Song, F., Astuti, G. D., Huynen, M. A., van Wijk, E., Stieger, K., & Collin, R. W. (2015). The pros and cons of vertebrate animal models for functional and therapeutic research on inherited retinal dystrophies. Progress in retinal and eye research, 48, 137–159. https://www.sciencedirect.com/science/article/pii/S1350946215000300?via%3Dihub

Smith, P. C., & Mong, J. A. (2019). Neuroendocrine control of sleep. Neuroendocrine Regulation of Behavior, 353–378.

Spiwoks-Becker, I., Maus, C., Tom Dieck, S., Fejtová, A., Engel, L., Wolloscheck, T., Wolfrum, U., Vollrath, L., & Spessert, R. (2008). Active zone proteins are dynamically associated with synaptic ribbons in rat pinealocytes. Cell and tissue research, 333(2), 185–195.

Staudt, N., Giger, F. A., Fielding, T., Hutt, J. A., Foucher, I., Snowden, V., Hellich, A., Kiecker, C., & Houart, C. (2019). Pineal progenitors originate from a non-neural territory limited by FGF signalling. Development, 146(22). 10.1242/dev.171405

Stemerdink, M., García-Bohórquez, B., Schellens, R., Garcia-Garcia, G., Van Wijk, E., & Millan, J. (2021). Genetics, pathogenesis and therapeutic developments for Usher syndrome type 2. Human Genetics, 1–22.

Takahashi, J. S., & Zatz, M. (1982). Regulation of circadian rhythmicity. Science, 217(4565), 1104–1111.

The UniProt Consortium. (2024). UniProt: the Universal Protein Knowledgebase in 2025. Nucleic acids research, 53(D1), D609–D617. 10.1093/nar/gkae1010

Toyama, R., Chen, X., Jhawar, N., Aamar, E., Epstein, J., Reany, N., Alon, S., Gothilf, Y., Klein, D. C., & Dawid, I. B. (2009). Transcriptome analysis of the zebrafish pineal gland. Developmental Dynamics, 238(7), 1813–1826. 10.1002/dvdy.21988

Ullah, F., Zeeshan Ali, M., Ahmad, S., Muzammal, M., Khan, S., Khan, J., & Ahmad Khan, M. (2025). Current updates on genetic spectrum of usher syndrome. Nucleosides Nucleotides Nucleic Acids, 44(5), 337–360. 10.1080/15257770.2024.2344194

van den Bos, R., Mes, W., Galligani, P., Heil, A., Zethof, J., Flik, G., & Gorissen, M. (2017). Further characterisation of differences between TL and AB zebrafish (Danio rerio): Gene expression, physiology and behaviour at day 5 of the larval stage. PloS one, 12(4), e0175420. 10.1371/journal.pone.0175420

van Faassen, M., Bischoff, R., & Kema, I. P. (2017). Relationship between plasma and salivary melatonin and cortisol investigated by LC-MS/MS. Clinical Chemistry and Laboratory Medicine (CCLM), 55(9), 1340–1348.

van Wijk, E., van der Zwaag, B., Peters, T., Zimmermann, U., Te Brinke, H., Kersten, F. F., Märker, T., Aller, E., Hoefsloot, L. H., Cremers, C. W., Cremers, F. P., Wolfrum, U., Knipper, M., Roepman, R., & Kremer, H. (2006). The DFNB31 gene product whirlin connects to the Usher protein network in the cochlea and retina by direct association with USH2A and VLGR1. Hum Mol Genet, 15(5), 751–765. 10.1093/hmg/ddi490

Vandesompele, J., De Preter, K., Pattyn, F., Poppe, B., Van Roy, N., De Paepe, A., & Speleman, F. (2002). Accurate normalization of real-time quantitative RT-PCR data by geometric averaging of multiple internal control genes. Genome biology, 3(7), 1–12.

Vatine, G., Vallone, D., Gothilf, Y., & Foulkes, N. S. (2011). It’s time to swim! Zebrafish and the circadian clock. FEBS letters, 585(10), 1485–1494. https://febs.onlinelibrary.wiley.com/doi/pdfdirect/10.1016/j.febslet.2011.04.007?download=true

Verbakel, S. K., van Huet, R. A. C., Boon, C. J. F., den Hollander, A. I., Collin, R. W. J., Klaver, C. C. W., Hoyng, C. B., Roepman, R., & Klevering, B. J. (2018). Non-syndromic retinitis pigmentosa. Progress in retinal and eye research, 66, 157–186. 10.1016/j.preteyeres.2018.03.005

Wahlqvist, M., Möller, C., Möller, K., & Danermark, B. (2013). Physical and psychological health in persons with deafblindness that is due to Usher syndrome Type II. Journal of Visual Impairment & Blindness, 107(3), 207–220.

Wang, M., Zhong, Z., Zhong, Y., Zhang, W., & Wang, H. (2015). The zebrafish period2 protein positively regulates the circadian clock through mediation of retinoic acid receptor (RAR)-related orphan receptor α (Rorα). Journal of Biological Chemistry, 290(7), 4367–4382. https://www.ncbi.nlm.nih.gov/pmc/articles/PMC4326843/pdf/zbc4367.pdf

Westerfield, M. (2007). The Zebrafish Book; A guide for the laboratory use of zebrafish (Danio rerio). *(No Title)*.

Yan, D., & Liu, X. Z. (2010). Genetics and pathological mechanisms of Usher syndrome. Journal of human genetics, 55(6), 327. https://www.ncbi.nlm.nih.gov/pmc/articles/PMC4511090/pdf/nihms707840.pdf

Yu, D., Zou, J., Chen, Q., Zhu, T., Sui, R., & Yang, J. (2020). Structural modeling, mutation analysis, and in vitro expression of usherin, a major protein in inherited retinal degeneration and hearing loss. Computational and Structural Biotechnology Journal, 18, 1363–1382. 10.1016/j.csbj.2020.05.025

Ziv, L., Tovin, A., Strasser, D., & Gothilf, Y. J. E. e. r. (2007). Spectral sensitivity of melatonin suppression in the zebrafish pineal gland. 84(1), 92–99.

Zrada, S. E., Braat, K., Doty, R. L., & Laties, A. M. (1996). Olfactory loss in Usher syndrome: another sensory deficit? American journal of medical genetics, 64(4), 602–603. https://onlinelibrary.wiley.com/doi/pdfdirect/10.1002/ajmg.1320640402?download=true

Bio-Atlas. (2013). Zebrafish atlas. https://bio-atlas.psu.edu/. Funding: NIH grant 5R24 RR01744, Jake Gittlen Cancer Research Foundation, and PA Tobacco Settlement Fund. Saggital section, 6dpf: http://bio-atlas.psu.edu/zf/view.php?s=35525&atlas=57

